# A tonoplast cytokinin riboside transporter gates intracellular hormone availability at the root–microbe interface

**DOI:** 10.64898/2026.05.31.727005

**Authors:** Martin Hudeček, Daniel Nedvěd, Anna Kuchařová, Sándor Tamás Forczek, Marharyta Kuzmenko, Petr Klíma, Vladimír Skalický, Rupali Gupta, Loida Tejada, Ljupka Trajkova Petrova, Olga Šamajová, Jozef Šamaj, Jakub Hajný, David Zalabák, Torsten Möhlmann, Maya Bar, Ondřej Novák, Eva Benková, Klára Hoyerová, Ondřej Plíhal

## Abstract

Cytokinin ribosides are major mobile and precursor forms of cytokinins, plant hormones whose transport and subcellular distribution shape developmental and stress responses. Here, we identify *Arabidopsis thaliana* EQUILIBRATIVE NUCLEOSIDE TRANSPORTER1 (ENT1) as a tonoplast-localized cytokinin riboside transporter. Tissue-specific subcellular analysis under native regulatory elements localized ENT1 predominantly to the tonoplast of root epidermal and lateral root cap cells, where it gates intracellular cytokinin riboside availability. Accordingly, *ENT1* overexpression enhanced cytokinin riboside sensitivity and signalling, whereas loss of *ENT1* altered adenosine metabolism and disrupted cytokinin homeostasis, leading to the accumulation of multiple zeatin-type cytokinins. ENT1-dependent cytokinin riboside compartmentalization was required for beneficial microbe-induced protection, as *ent1* mutants failed to acquire protection against the fungal pathogen *Botrytis cinerea* and the bacterial pathogen *Pseudomonas syringae* pv. tomato DC3000. These findings reveal a vacuolar gatekeeping mechanism that controls intracellular cytokinin riboside availability and links hormone compartmentalization to beneficial microbe-dependent plant defence.

## INTRODUCTION

Cytokinins (CKs) coordinate multiple developmental and stress-related processes throughout the plant life cycle, including root and shoot meristem activity, organ formation and broader developmental patterning^1^, photomorphogenic development and photosynthetic regulation^2,3^, and defence responses^4^.

CKs occur in multiple molecular forms, including free bases, ribosides, nucleotides, and glucosides. Among these, free bases are the primary ligands of CK receptors^5^, membrane-associated hybrid histidine kinases (HKs) containing a CHASE ligand-binding domain^6,7^. By contrast, CK ribosides generally show much lower receptor affinity and are therefore considered primarily transport and precursor forms rather than principal signalling species^5,8^.

CK output is further shaped by the metabolic activation of CK riboside 5′-monophosphates by LONELY GUY (LOG) phosphoribohydrolases, which directly generate bioactive free bases^9,10^. Conversely, CK signalling is attenuated by irreversible cytokinin oxidase/dehydrogenase (CKX)-mediated degradation^11^ and by UDP-glycosyltransferase (UGT)-mediated glucosylation, mainly at the N7 and N9 positions of the adenine moiety or at the O-position of the zeatin side chain^12–14^. O-glycosylated CKs are not biologically active but can be reactivated to free bases by β-glucosidases^15^.

The cellular and subcellular partitioning of active, metabolic, transport, or inactivated CK forms may influence where and when active CK pools arise. Transport is therefore one of the major control points in CK biology. Root-derived *trans-*zeatin (*t*Z)-type CKs, including *trans*-zeatin riboside (*t*ZR), contribute to long-distance root-to-shoot communication, particularly in nutrient-responsive growth regulation^16,17^. Several CK transport components have been identified, including root-to-shoot *t*Z-type transporter ATP-binding cassette G14 (ABCG14)^18,19^, and AZA-GUANINE RESISTANT 1 and 2 (AZG1/2), which mediate developmental responses in roots in concert with auxin^20,21^. However, the transport processes that control CK distribution at the cellular and subcellular levels remain incompletely understood.

Members of the evolutionarily ancient family of EQUILIBRATIVE NUCLEOSIDE TRANSPORTER (ENT) are strong candidates for CK riboside transport. In Arabidopsis, eight members have been identified^22–24^, and several family members have already been linked to CK riboside transport^25–28^. ENT6 transports CK ribosides in a heterologous system^26^, and recent work demonstrated that ENT3 mediates both *t*ZR and isopentenyladenosine (iPR) transport in a plant cell-based system^28^. By contrast, the contribution of ENT1 to CK transport has remained unresolved, despite its established function as a nucleoside transporter associated with vacuolar nucleoside export^29,30^.

Given its tonoplast localization and nucleoside transport activity, ENT1 is a strong candidate for mediating the retrieval of CK ribosides from the vacuole into the cytosol. This step remains poorly understood, although vacuoles contain biologically relevant pools of active, storage, and transport-associated CK forms^31–34^. Defining whether ENT1 transports CK ribosides is therefore important for understanding how intracellular CK partitioning regulates local hormone availability and downstream CK signalling outputs.

Intracellular CK partitioning may also be relevant at plant–microbe interfaces. In tomato, altered CK content or CK sensitivity reshapes the phyllosphere microbiome and contributes to microbiome-dependent pathogen resistance^35^. Interestingly, *Trichoderma* species produce CK pools enriched in riboside and nucleotide derivatives^36^, and beneficial *Bacillus* isolates can activate CK-responsive genes in the plant host, including signalling pathway components and *CKX* genes, together with defence-associated markers^37^. A more direct functional link was shown for *Pseudomonas fluorescens* G20-18, whose CK-deficient bacterial mutants displayed impaired biocontrol against the plant pathogen *Pseudomonas syringae* in Arabidopsis^38^. Whether CK transport contributes to this layer of plant-microbe communication remains unknown.

Here, we show for the first time that ENT1 functions as a selective CK riboside transporter, define its substrate specificity, and confirm its subcellular localization through live-cell imaging. We further demonstrate that ENT1-dependent intracellular transport reshapes endogenous CK riboside pools, thereby modulating CK responsiveness, beneficial microbe-mediated immune priming, and pathogen protection *in planta*.

## RESULTS

### ENT1 is a transporter of cytokinin ribosides but not nucleobases

To determine whether ENT1 transports CKs, we generated a BY-2 cell line expressing an inducible ENT1-eGFP under the control of the β-estradiol-inducible *XVE* transactivation system^39,40^, hereafter referred to as *XVE>>ENT1^WT^-eGFP.* CK uptake was subsequently quantified by radioaccumulation assays using CK nucleobases and ribosides.

Induction of *ENT1* expression by β-estradiol increased the first-order uptake rate constants for [2-³H]*t*ZR and [2-³H]iPR compared with non-induced control cells. In contrast, *ENT1* induction did not affect uptake of the CK nucleobases [2-³H]*t*Z and [2-³H]iP, indicating that ENT1 preferentially transports CK ribosides (Fig. 1a, Supplementary Fig. 1a).

**Fig. 1.**
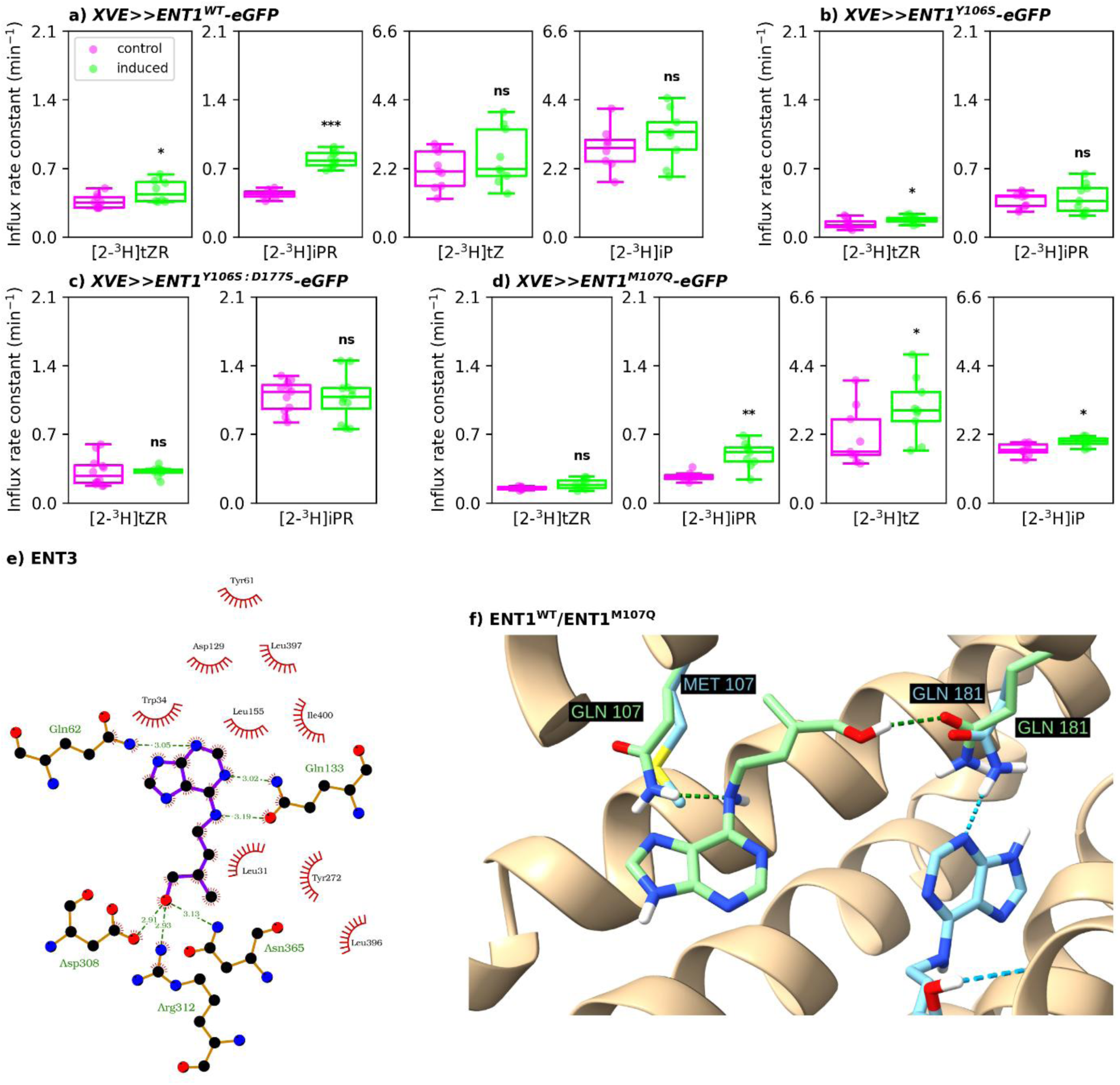
ENT1 selectively transports cytokinin ribosides and its substrate specificity depends on conserved binding-site residues. **a–d**, Radioaccumulation assays in BY-2 cell suspensions expressing β-estradiol-inducible *ENT1* variants. Uptake of [2-^3^H]*t*ZR, [2-^3^H]iPR, [2-^3^H]*t*Z, and [2-^3^H]iP was measured in **a**, *XVE>>ENT1^WT^-eGFP*; **b**, *XVE>>ENT1^Y^*^106^*^S^-eGFP*; **c**, *XVE>>ENT1^Y^*^106^*^S:D^*^177^*^S^-eGFP*; and **d**, *XVE>>ENT1^M^*^107^*^Q^-eGFP*. The first-order uptake rate constant was calculated from parameters obtained by fitting Eq. (1) to time-course data of intracellular radioactivity. **e**, Interaction scheme for the best-scoring docked pose of *t*Z in the AlphaFold-predicted ENT3 of *A. thaliana*. Dashed lines denote hydrogen bonds and short concentric lines denote apolar interactions. Visualized in LigPlot+. **f**, Best-scoring *t*Z poses obtained by molecular docking into AlphaFold-predicted models of wild-type ENT1 and ENT1^M107Q^. Dashed lines denote hydrogen bonds. Structures were visualized in ChimeraX. Boxplots show the median and interquartile range (IQR); whiskers extend to the most extreme non-outlier values within 1.5 × IQR, and points indicate individual measurements. Statistical significance of differences between induced and control suspensions was assessed using a Kruskal-Wallis test; *\*P* < 0.05, *\*\*P* < 0.01, *\*\*\*P* < 0.001; ns, not significant.

To investigate whether Tyr106 and Asp177 contribute to CK substrate recognition, as shown previously for the corresponding residues in ENT3^28^, we generated *XVE>>ENT1^Y^*^106^*^S^-eGFP* and *XVE>>ENT1^Y^*^106^*^S:D^*^177^*^S^-eGFP* constructs and introduced them into BY-2 cells. Upon induction, neither substituted ENT1 variant increased the uptake of [2-³H]*t*ZR or [2-³H]iPR, unlike the WT construct, supporting a role for Tyr106 and Asp177 in CK substrate binding (Fig. 1b,c, Supplementary Fig. 1b,c).

Unlike ENT1, ENT3 transports CK nucleobases in BY-2 cells^28^. To identify residues that may contribute to this difference in substrate specificity, we docked *t*Z into an AlphaFold-generated model of ENT3 (Fig. 1e) and assessed the conservation of residues predicted to interact with the ligand (Supplementary Fig. 1e). This analysis identified Gln62, a residue predicted to interact with *t*Z in ENT3, as corresponding to Met107 in ENT1. We therefore hypothesized that this non-conserved binding-site residue contributes to the distinct CK nucleobase affinity of ENT1 and ENT3. To test this hypothesis, we next docked *t*Z into AlphaFold-generated models of wild-type ENT1 (ENT1^WT^) and M107Q-substituted ENT1 (ENT1^M107Q^). In the ENT1^M107Q^ model, Gln107 formed a hydrogen bond with *t*Z, altering the predicted ligand-binding pose (Fig. 1f). To experimentally test this prediction, we generated the *XVE>>ENT1^M^*^107^*^Q^-eGFP* construct and introduced it into BY-2 cells. Radioaccumulation assays showed that ENT1^M107Q^ increased uptake of *t*Z and iP, unlike ENT1^WT^. Notably, ENT1^M107Q^ retained transport activity towards iPR, whereas transport of *t*ZR was reduced (Fig. 1d, Supplementary Fig. 1d).

### ENT1 exhibits a distinct epidermal expression pattern in roots and aerial tissues

To refine the *in planta* expression of *ENT1* at cellular resolution, we generated an independent dual reporter line expressing a nuclear-targeted eGFP-GUS fusion under the control of a 4 kb *ENT1* upstream region and an 840 bp downstream region (*pENT1::NLS-eGFP-GUS:tENT1*), expanding the regulatory context relative to the previously used 1.1 kb promoter-only reporter^30^.

The reporter expression strongly marked epidermal cells along the entire root axis, including the root apical meristem (RAM) (Fig. 2a). In the RAM, epidermal expression was predominantly enriched in atrichoblasts (Fig. 2a), lateral root cap cells (LRC) (Fig. 2b,c), and in the root hairs within the differentiation zone (Fig. 2d).

**Fig. 2.**
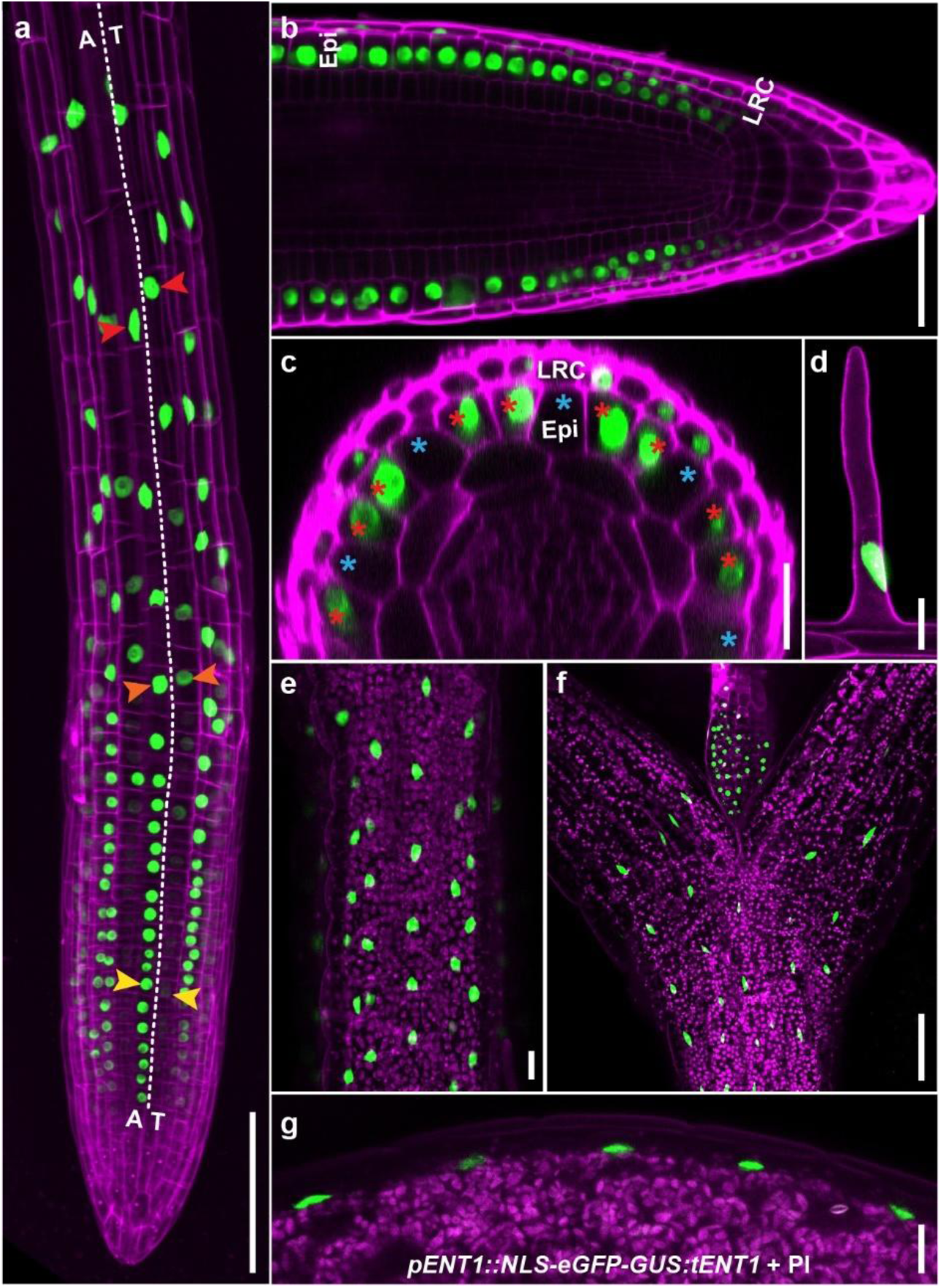
ENT1 reporter activity is restricted to epidermal and lateral root cap tissues. **a-g**, Co-imaging of *pENT1::NLS-eGFP-GUS:tENT1* reporter activity with propidium iodide (PI) labelling of cell walls in living cells (provided in false magenta colour, **a-d**) and/or chlorophyll autofluorescence in plant green organs and tissues (**e-g**) using laser confocal microscopy. **a**, Root of a 3-day-old seedling. Yellow, orange, and red arrowheads indicate nuclei in neighbouring atrichoblast (A) and trichoblast (T) cell files in the meristematic, transition, and elongation root zones, respectively. The dashed line marks the boundary between the adjacent A and T epidermal cell files and serves as a guide to compare reporter signal along the root axis. Difference in the reporter intensity between A and T cell files progressively diminished in the elongation zone (red arrowheads). **b**, Longitudinal view of the root cap and meristematic zone of a 3-day-old seedling. **c**, Transverse section of the root meristematic zone covered by lateral root cap cells (LRC) of a 3-day-old seedling. Red and blue asterisks indicate A and T cells, respectively. **d**, Root hair of a 7-day-old seedling. **e**, Hypocotyl of a 3-day-old seedling. **f**, Upper hypocotyl, cotyledon bases, and emerging first true leaf of a 7-day-old seedling. **g**, Cotyledon margin in a 3-day-old seedling. Epi, epidermis; LRC, lateral root cap; PI, propidium iodide. Scale bars: 100 μm (**a**,**f**), 50 μm (**b**), 20 μm (**c–e**,**g**).

Reporter activity was also detected in the aerial epidermal tissues of developing Arabidopsis seedlings (Fig. 2e-g). Signal was observed in hypocotyl epidermal cells of 3-day-old seedlings and extended to the epidermis of the first true leaves in 7-day-old seedlings (Fig. 2e,f). In addition, reporter activity was detected in cotyledons (Fig. 2g).

Using a reporter with high cellular resolution, we refined the previously described broad root expression pattern of *ENT1*^30^ and revealed a specific epidermis-enriched localization.

### Internal tagging reveals tonoplast localization of ENT1 and an overexpression-driven shift to the plasma membrane

Previous studies reported conflicting evidence for ENT1 localization, with plasma membrane (PM) targeting suggested in a heterologous system and tonoplast localization supported by vacuolar proteomics and immunodetection^30,41,42^.

To determine ENT1 localization *in planta* by live-cell imaging, we generated stable transgenic Col-8 lines producing internally tagged ENT1 fusion proteins carrying either the wild-type sequence (ENT1^WT^-eGFP) or a targeted mutation in the putative acidic di-leucine motif (Supplementary Fig. 2a). As this motif is a candidate tonoplast-sorting signal, we substituted Leu45 with glycine to test whether it contributes to ENT1 tonoplast targeting, generating ENT1^L45G^-eGFP. These constructs were expressed in the Col-8 background under native *ENT1* 5′ and 3′ regulatory regions, providing a near-native regulatory context but elevated expression relative to the endogenous locus (Fig. 3a).

**Fig. 3.**
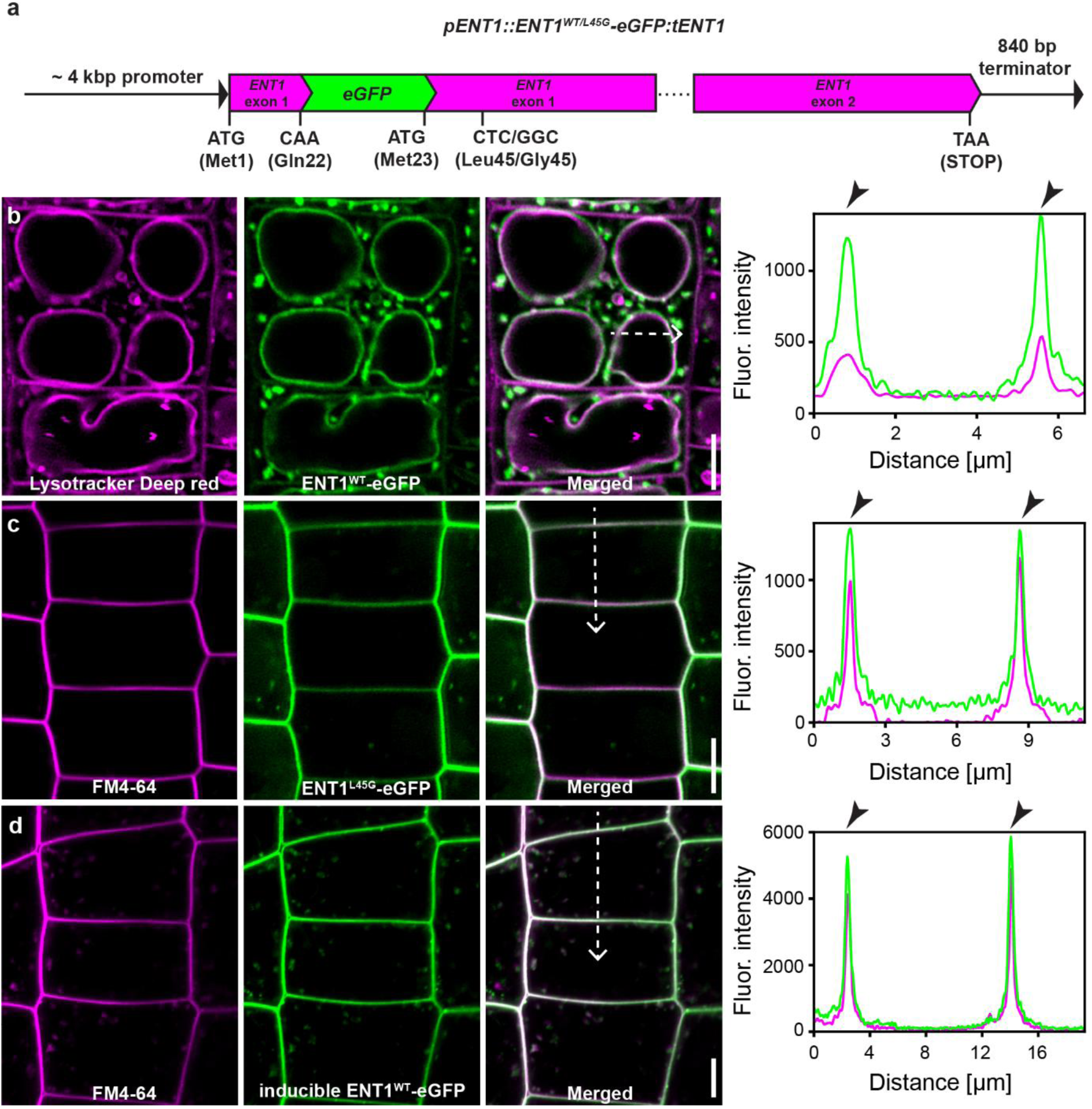
Tonoplast localization of ENT1 depends on its N-terminal sorting motif and expression context. **a**, Schematic presentation of the internally tagged *pENT1::ENT1^WT^-eGFP:tENT1* and *pENT1::ENT1^L45G^-eGFP:tENT1* constructs, each containing a 4 kb *ENT1* 5′ upstream region, the *ENT1* genomic sequence with *eGFP* inserted between Gln22 and Met23, and an 840 bp 3′ downstream region, both introduced in *A. thaliana* Col-8 background. **b–d**, Imaging of root epidermal cells of 3-day-old *A. thaliana* Col-8 seedlings using laser confocal microscopy. **b**, Fusion protein ENT1^WT^-eGFP co-localized with the tonoplast marker LysoTracker Deep Red. **c**, Mutated fusion protein ENT1^L45G^-eGFP accumulated at the plasma membrane (PM), as indicated by co-localization with PM vital stain FM4-64. **d**, β-estradiol-inducible ENT1^WT^-eGFP also accumulated at the PM following strong induction. In all panels, eGFP fluorescence is shown in green, and tonoplasts and PM markers are depicted in magenta. Fluorescence intensity profiles in both channels were measured along the white dashed arrows, starting from the upper end (0 µm), as indicated in **b–d**; black arrowheads on the right indicate overlapping fluorescence maxima. Scale bars, 5 µm.

Because acidic di-leucine motifs can mediate tonoplast targeting of plant membrane proteins^43,44^, we inspected the conservation of the ENT1 ETSLLL (amino acid residues 41-46) motif across the Arabidopsis ENT family. The aligned Arabidopsis ENT sequences showed that this motif is not conserved among other Arabidopsis ENT paralogues, but was specific to ENT1 (Supplementary Fig. 2b) and may affect its localization (Supplementary Fig. 2c).

In ENT1^WT^-eGFP seedlings, confocal imaging showed that this fusion protein predominantly localized to the vacuolar membrane in root epidermal cells, and co-localized with LysoTracker Deep Red at the vacuolar boundary, consistent with tonoplast localization (Fig. 3b). Contrary to this, live-cell imaging of ENT1^L45G^-eGFP seedlings showed predominant PM localization accompanied by co-localization with FM4-64, whereas tonoplast-associated fluorescent signal was strongly reduced or absent (Fig. 3c).

PM localization was also observed after strong β-estradiol induction of *XVE*>>*ENT1^WT^-eGFP* expression (36.89-fold; Supplementary Fig. 3c), despite the intact ETSLLL motif (Fig. 3d).

Together, these findings suggest that efficient tonoplast localization of ENT1 depends on both the integrity of the putative acidic di-leucine motif and the expression context, as both ETSLLL motif disruption and strong inducible expression were associated with increased PM accumulation.

### Elevated ENT1 abundance promotes sensitivity to cytokinin ribosides

Since CK ribosides, particularly *t*ZR, are important mobile forms in vascular CK distribution^16,45^, we tested whether altered ENT1 abundance affects plant sensitivity to CK ribosides *in planta*.

We first characterized two available Arabidopsis *ent1* T-DNA insertion alleles, *ent1-1* and *ent1-2*, each carrying an insertion in the first exon (Supplementary Fig. 4a-f). Both lines retained residual *ENT1* transcript variants and low, but still detectable ENT1 protein levels (Supplementary Fig. 4g). We therefore refer to them hereafter as *ent1* knock-down lines.

Next, we generated two independent CRISPR-Cas9 mutant lines, *ent1-3* and *ent1-4* (Supplementary Fig. 5). Both alleles are predicted to disrupt the *ENT1* coding sequence and introduce a premature termination codon. Consequently, ENT1 protein was not detectable by immunoblotting with an anti-ENT1 antibody in both CRISPR alleles (Supplementary Fig. 4g). Nevertheless, *ent1-4* did not display a discernible phenotype under standard growth conditions, or when grown *in vitro* in the presence of CK ribosides *t*ZR and iPR (Supplementary Fig. 6), indicating that loss of *ENT1* is not essential for gross whole-seedling responses to exogenously supplied CK ribosides under these conditions.

We therefore used complementary overexpression approaches to test whether increased *ENT1* abundance affects plant growth or sensitivity to CK ribosides. For this purpose, we analysed two mildly overexpressing lines generated under *ENT1* 5′ and 3′ regulatory regions: predominantly tonoplast-localized ENT1^WT^-eGFP and PM-localized ENT1^L45G^-eGFP, and tested their responses to 100 nM iPR or *t*ZR treatments compared to Col-8 (Fig. 4a,d). *ENT1* transcript levels were mildly increased relative to Col-8, with 1.64-fold higher expression in ENT1^WT^-eGFP (Fig. 4c) and 2.87-fold higher expression in ENT1^L45G^-eGFP (Fig. 4f).

**Fig. 4.**
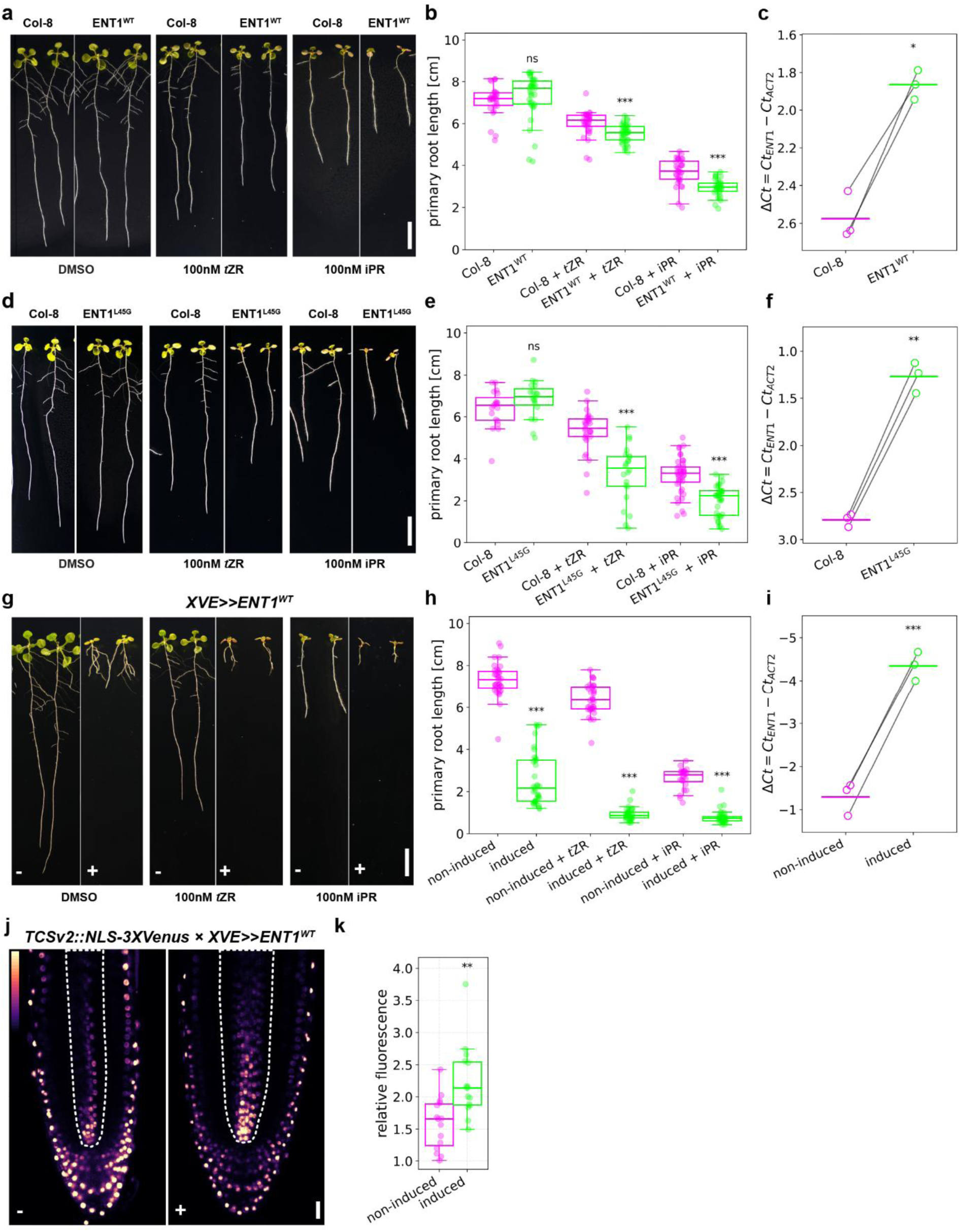
Increased ENT1 abundance enhances sensitivity to cytokinin ribosides and elevates cytokinin signalling output. **a**, Representative phenotypes of 10-day-old *A. thaliana* Col-8 and ENT1^WT^-eGFP line expressing *ENT1^WT^-eGFP* under the native *ENT1* regulatory context. Seedlings were grown on control medium containing DMSO or the same medium supplemented with 100 nM *t*ZR or 100 nM iPR. **b**, Primary root length for the genotypes and conditions as shown in **a**. **c**, RT-qPCR analysis of *ENT1* transcript levels in *A. thaliana* Col-8 and ENT1^WT^-eGFP seedlings. Expression is plotted as Δ*Ct* = *Ct_ENT1_* − *Ct_ACT2_*; the y-axis is inverted, so lower Δ*Ct* values indicate higher expression. Open circles represent independent experiments (*n* = 3), paired experiments are connected by lines, and bars indicate means. **d–f,** As in **a–c**, but for *A. thaliana* Col-8 and ENT1^L45G^-eGFP seedlings expressing *ENT1^L45G^-eGFP* under the native *ENT1* regulatory context. **g**, Representative phenotypes of 10-day-old XVE>>ENT1^WT^ seedlings kept non-induced (−) or induced with 5 μM β-estradiol (+) on medium without cytokinins (DMSO) or supplemented with 100 nM *t*ZR or 100 nM iPR. **h**, Primary root length of seedlings shown in **g**. **i**, RT-qPCR of *ENT1* transcript levels in XVE>>ENT1^WT^ seedlings kept non-induced or induced with 5 μM β-estradiol, presented as in **c**. **j**, Representative confocal images of *TCSv2::NLS-3XVenus* × *XVE>>ENT1^WT^* in root apices of 7-day-old seedlings kept non-induced (−) or induced with 10 µM β-estradiol (+). Dashed outlines indicate the stele region used for quantification. Pseudocolour indicates Venus fluorescence intensity. **k**, Mean Venus fluorescence intensity in the root stele of seedlings shown in **j**; points represent individual seedlings (*n* = 15). Boxplots in **b**, **e**, **h**, and **k** show the median and interquartile range (IQR); whiskers extend to the most extreme non-outlier values within 1.5 × IQR, and points indicate individual seedlings. Statistical significance was assessed using Welch’s two-sample t-test (**b**, **e**, **h**), or a two-tailed Mann–Whitney U test (**k**). Statistical significance in **c**, **f,** and **i** was assessed using a paired t-test. *\*P* < 0.05, *\*\*P* < 0.01, *\*\*\*P* < 0.001; ns, not significant. Scale bars, 1 cm (**a**, **d**, **g**), and 20 µm (**j**).

Both lines displayed increased CK sensitivity relative to Col-8 (Fig. 4, Supplementary Table 2), consistent with CK hypersensitivity phenotypes previously reported in *IPT*-overexpressing lines^46^, including reduced root meristem size and suppressed lateral root formation. Both ENT1^WT^-eGFP and ENT1^L45G^-eGFP showed stronger primary root growth inhibition on both *t*ZR and iPR compared with Col-8 (Fig. 4b,e). By contrast, treatment with the active free-base CKs iP and *t*Z at 50 nM caused comparable growth inhibition in both genotypes, indicating that observed changes were specifically caused by CK ribosides (Supplementary Fig. 7).

We also generated β-estradiol-inducible *XVE>>ENT1^WT^*Arabidopsis lines strongly overexpressing untagged *ENT1^WT^*. Upon induction, *ENT1^WT^* overexpression caused a pronounced root growth defect even on mock medium (Fig. 4g-i). CK hypersensitivity can be evident even under mock conditions, likely reflecting enhanced responsiveness to endogenous CK pools^47^. This effect was further enhanced with exogenous CK riboside supplementation, as induction of *ENT1^WT^* led to a markedly stronger reduction in median primary root length (Fig. 4h). To further support this, we tested eGFP-tagged, β-estradiol-inducible *XVE>>ENT1^WT^-eGFP*-expressing line, which likewise displayed a pronounced CK riboside hypersensitivity phenotype (Supplementary Fig. 3a,b). The corresponding increase in ENT1 protein abundance in individual lines was confirmed by immunoblotting with a custom-made ENT1-specific antibody (Supplementary Fig. 8a,b).

To test whether the root growth phenotypes in the untagged inducible line were associated with enhanced CK signalling, we crossed *XVE>>ENT1^WT^*-expressing line with the reporter line carrying *TCSv2::NLS-3XVenus* construct^48^. Upon induction with β-estradiol, Venus fluorescence increased significantly in the root stele relative to the non-induced control (Fig. 4j,k).

Overall, these results implicate CK ribosides as preferred ENT1 substrates *in planta*. Increased sensitivity under both native-promoter expression and strong inducible expression was specific to ribosylated CK forms. In contrast, responses to free-base CKs remained largely unchanged, suggesting that their transport is mediated by other transport systems.

### Altered ENT1 activity shapes cytokinin riboside homeostasis

Although the *ent1-4* null mutant did not show a pronounced phenotype under standard growth conditions or an altered growth response to exogenously supplied CK ribosides, the tonoplast localization of ENT1 suggests that absent ENT1 function could still reshape CK metabolite fluxes. We therefore quantified CK species and adenosine (Ado) in roots and shoots of 12-day-old Col-8 and *ent1-4* seedlings grown on either mock medium or medium supplemented with 100 nM *t*ZR.

Loss of *ENT1* caused the strongest CK homeostasis perturbation in mock-grown roots, where multiple active CKs, CK ribosides, and riboside-related metabolites accumulated (Fig. 5a, Supplementary Fig. 9). These included *c*Z and iP among free-base CKs, all major CK ribosides analysed, and the O-glucosylated *c*ZR derivative *c*ZROG. Ado was also increased in mock-grown *ent1-4* roots, consistent with the established role of ENT1 in Ado transport and/or altered CK riboside turnover (Fig. 5a). By contrast, *t*ZR-grown *ent1-4* roots showed a more restricted shift, mainly involving increased *t*ZROG, *c*ZR and *c*ZROG and reduced *c*ZRMP (Fig. 5a, Supplementary Fig. 9).

**Fig. 5.**
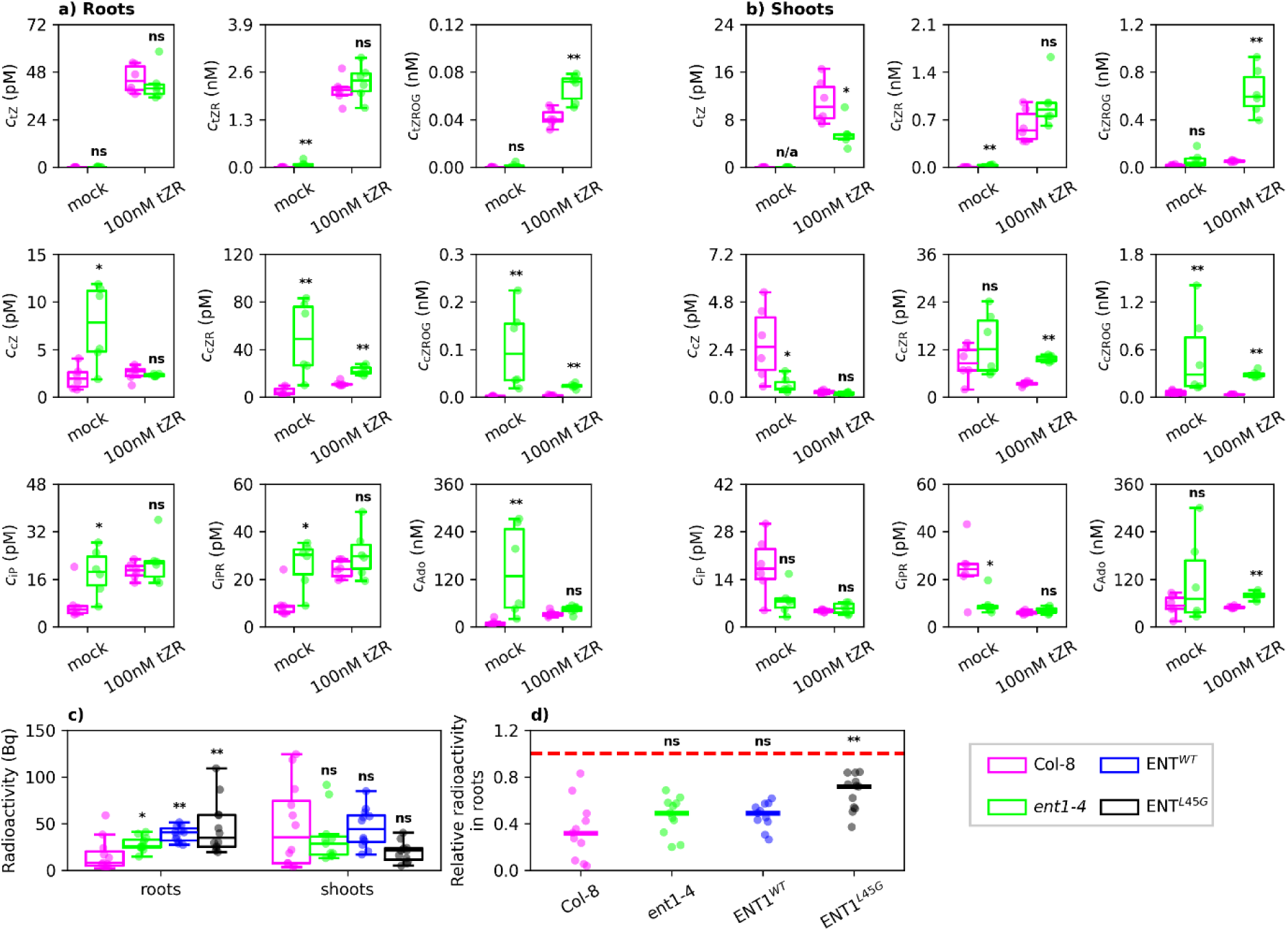
ENT1 activity modulates cytokinin riboside homeostasis in roots and shoots of *A. thaliana* seedlings. Cytokinin-related metabolite profiles in **a**, roots and **b**, shoots of 12-day-old *A. thaliana* Col-8 and *ent1-4* plants treated with DMSO mock or 100 nM *t*ZR. Shown are *t*Z, *t*ZR, *t*ZROG, *c*Z, *c*ZR, *c*ZROG, iP, iPR, and adenosine (Ado) levels. Notable changes are visible in *c*ZR, *c*ZROG, Ado, and iPR levels in *ent1-4* mock-grown roots. **c**, Radioactivity measured in roots and shoots of 8-day-old *A. thaliana* seedlings of indicated genotypes after 4 h treatment of root tips with 79.21 MBq l^−1^ [2-^3^H]*t*ZR. **d**, root radioactivity from **c** expressed relative to total radioactivity in whole plants. The dashed line is plotted at y = 1.0. Boxplots show the median and interquartile range (IQR); whiskers extend to the most extreme non-outlier values within 1.5 × IQR, and points indicate individual measurements. Short horizontal lines in **d** show medians. Statistical significance between wild-type Col-8 and the indicated genotypes was assessed using a Kruskal-Wallis test; *\*P* < 0.05, *\*\*P* < 0.01, *\*\*\*P* < 0.001; ns, not significant.

*ENT1* loss altered CK profiles in shoots, but in a more selective pattern than in roots. In mock-grown *ent1-4* shoots, while *c*Z was reduced, *t*ZR and *c*ZROG increased and iPR decreased (Fig. 5b, Supplementary Fig. 9). In *t*ZR-grown plants, *ent1-4* shoots again accumulated *c*ZR and *c*ZROG, accompanied by reduced *t*Z and an increased Ado level (Fig. 5b, Supplementary Fig. 9). Together, these profiles indicate that *ENT1* loss reconfigures riboside-related CK pools in a tissue- and treatment-dependent manner, with *c*ZROG consistently elevated across both tissues and treatments.

Because altered *t*ZR and *t*ZROG levels could reflect changes in *t*ZR-related root-to-shoot distribution, we applied radiolabelled [2-³H]*t*ZR to root tips of Col-8, *ent1-4*, ENT1^WT^-eGFP, and ENT1^L45G^-eGFP seedlings. Root radioactivity was higher in all genotypes with altered ENT1 activity compared to Col-8, whereas shoot radioactivity did not differ significantly between genotypes (Fig. 5c). The proportion of *t*ZR-derived radioactivity retained in roots increased significantly only in ENT1^L45G^-eGFP seedlings (Fig. 5d). These data suggest that altered ENT1 activity affects root retention or redistribution of *t*ZR-derived compounds rather than bulk shootward delivery under these conditions.

To test whether altered CK riboside compartmentalization affects a CK-dependent developmental process, we used a hypocotyl explant regeneration assay^49,50^, again with *t*ZR as the CK source. Compared with Col-8, ENT1^L45G^-eGFP explants produced larger regenerated tissues, whereas the regenerated area did not differ significantly in *ent1-4* or ENT1^WT^-eGFP explants (Supplementary Fig. 10). Metabolite profiling of regenerated explants revealed genotype-dependent shifts in the same riboside-related metabolite groups affected in seedlings (Supplementary Fig. 10, 11). In *ent1-4*, *t*ZR, *t*ZROG, *c*ZR, and *c*ZROG accumulated, whereas ENT1^WT^-eGFP showed a broadly opposite pattern, with reduced *t*ZR, *c*ZR and *c*ZROG levels. ENT1^L45G^-eGFP explants accumulated *t*ZR, *t*ZROG, *c*ZROG, and iPR, and was the only line in which active CK free bases also accumulated (*t*Z and DHZ).

Together, these data show that ENT1 activity shapes CK homeostasis across tissues and developmental contexts. Rather than controlling bulk exogenous CK riboside responsiveness, ENT1 appears to regulate the intracellular partitioning and metabolic routing of riboside-derived CK pools. The limited effect of *t*ZR treatment, especially on free CK base pools in *ent1-4*, may help explain why this mutant does not show a gross growth phenotype upon exposure to exogenous CK ribosides, despite pronounced changes in riboside-related and O-glucosylated CK metabolites.

### ENT1 contributes to beneficial microbe-dependent protection in Arabidopsis

Loss of *ENT1* altered the homeostasis of multiple CK riboside-related metabolites in a tissue-specific manner. Given that *ENT1* is expressed in outer root tissues, we asked whether ENT1-dependent CK riboside compartmentalization contributes to plant responses to beneficial microbes, which are known to affect CK signalling outputs of the plant host ^36,37^. As modulation of root system architecture (RSA) is a hallmark of beneficial plant-microbe interactions^51,52^, we first examined whether the selected microbial strains altered root development.

Col-8 plants exhibited significant developmental responses to microbial treatments, including primary root growth inhibition and increased root hair length after treatment with *T. harzianum* T39, *B. megaterium* 4C, and *B. pumilus* R2E isolates (Supplementary Fig. 12a,b). These root responses were absent in *ent1-4* and attenuated in *ent1-2*, suggesting that ENT1-mediated intracellular CK partitioning contributes to the developmental competence of roots to respond to beneficial microbial cues (Supplementary Fig. 12a,b).

Because CK signalling has been linked to pathogen resistance and beneficial microbe-mediated protection^38,53,54^, we next tested whether ENT1-mediated intracellular CK distribution contributes to microbe-induced protection against common pathogens, *Botrytis cinerea* and *Pseudomonas syringae* pv. tomato DC3000. In Col-8, T39, 4C, and R2E pretreatments reduced grey mould disease caused by *Botrytis cinerea* (Fig. 6a). This effect was attenuated in the T-DNA insertion lines *ent1-1* and *ent1-2*, and a similar attenuation was observed in the independent CRISPR-Cas9 line *ent1-4* (Fig. 6a). In Col-8, the same microbes also reduced *Pseudomonas syringae* pv. tomato DC3000 proliferation (Fig. 6b) and water-soaked lesion development (Fig. 6c), whereas these protective effects were largely lost in *ent1-2* plants (Fig. 6b,c).

**Fig. 6.**
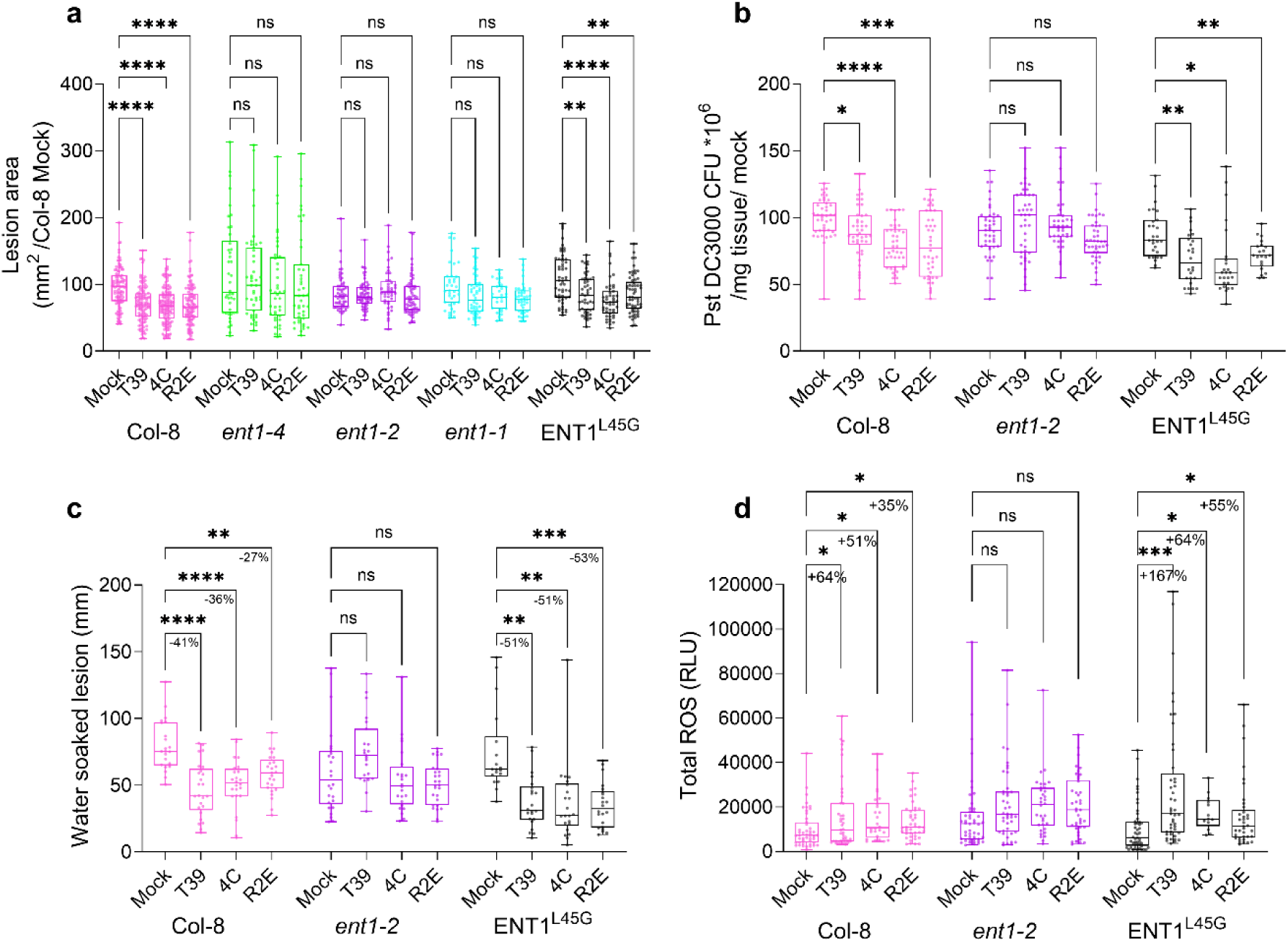
Beneficial microbe-induced disease protection requires functional cytokinin riboside homeostasis. **a–c**, Biocontrol activity of *Trichoderma harzianum* T39, *Bacillus megaterium* 4C, and *Bacillus pumilus* R2E was assessed in different *A. thaliana* Col-8 genotypes; 4-week-old plants were used for beneficial microbe pretreatments. Plants were treated with the indicated beneficial microbes (abbreviated T39, 4C, or R2E) 2 times: 4 days and 1 day before pathogen infection. **a**, Grey mould lesion area 3 days after inoculation of detached mature rosette leaves with *Botrytis cinerea* mycelia. **b**, *Pseudomonas syringae* pv. *tomato* DC3000 proliferation, measured as colony-forming units, 3 days after syringe infiltration of mature rosette leaves. **c**, Water-soaked lesion area caused by *P. syringae* pv. *tomato* DC3000 at 3 days post-inoculation. Protective effects of beneficial microbe pretreatments in panels **a–c** were statistically significant in Col-8 and ENT1^L45G^-eGFP lines incited by pathogens, whereas *ent1* knock-down and knock-out lines did not show significant protection under these conditions. **d**, flg22-induced reactive oxygen species (ROS) burst in leaf discs from the indicated genotypes after beneficial microbe pretreatment. ROS production was measured immediately after application of 1 µM flg22, with measurements taken at 3-min intervals for 40 min using the luminol–peroxidase assay and expressed as total relative luminescence units (RLU). Boxplots show the median and interquartile range; whiskers indicate the full data range, and points represent individual biological measurements from at least three independent experiments. Statistical significance indicates disease reduction (**a–c**) or increased ROS production (**d**) relative to the corresponding mock treatment within each genotype, assessed using Welch’s ANOVA followed by Dunnett’s post hoc test (**a–c**) or Kruskal–Wallis ANOVA followed by Dunn’s post hoc test (**d**). Sample sizes: **a**, *n* > 40; **b**, *n* > 25; **c**, *n* > 18; **d**, *n* > 15. *\*P* < 0.05, *\*\*P* < 0.01, *\*\*\*P* < 0.001, *\*\*\*\*P* < 0.0001; ns, not significant.

As beneficial microbes can potentiate early pattern-triggered immune (PTI) responses^55^, we next quantified the flg22-induced reactive oxygen species (ROS) burst after microbial pretreatment. This assay is relevant to CK-associated immunity because CK signalling has been linked to PAMP-triggered ROS production and stomatal defence against *Pseudomonas syringae*^56^. In Col-8, pretreatment with T39, 4C, or R2E enhanced flg22-triggered ROS production, whereas this response was not significantly increased in *ent1-2* plants (Fig. 6d).

Together, these results support a model in which ENT1-dependent intracellular partitioning of CK riboside-related metabolites contributes to beneficial microbe-induced RSA modulation, immune priming, and pathogen protection.

## DISCUSSION

Although ENT1 was characterized more than two decades ago as a nucleoside transporter^29,41^, its contribution to CK transport remained unresolved^23,25^. Here, we identify ENT1 as a tonoplast-localized CK riboside transporter that controls the intracellular availability of riboside-derived CK pools.

Transport assays, structural modelling and mutational analysis show that ENT1 preferentially transports CK ribosides, including *t*ZR and iPR, but not the corresponding free bases (Fig. 1). This riboside selectivity distinguishes ENT1 from broader-specificity ENT family members^28^ and is supported by residues that mediate riboside recognition and nucleobase exclusion.

Our work also resolves a long-standing uncertainty concerning ENT1 localization. Using live-cell imaging, we confirmed that ENT1 localizes predominantly to the tonoplast (Fig. 3). The L45G substitution redirected ENT1 to the PM, demonstrating that the acidic di-leucine motif, apparently unique to ENT1 among Arabidopsis ENTs, is required for tonoplast targeting. ENT1 is therefore specialized for intracellular transport at the tonoplast, where it can control the retrieval or redistribution of nucleosides and CK ribosides from vacuolar pools.

The dual localization of CK receptors at the PM^57,58^ and ER^59,60^ also highlights the importance of intracellular CK distribution, yet the transport processes that partition CK metabolites between intracellular compartments remain poorly understood. Vacuoles contain diverse CK forms, including active, storage, and transport-associated metabolites^31–34^. ENT1 therefore provides a molecular route by which CK riboside pools could be remobilized for cytosolic metabolism and signalling.

Consistent with this, loss of *ENT1* reconfigured CK homeostasis in a tissue-specific manner (Fig. 5). The most consistent effects involved *c*Z-type ribosides and their O-glucosides, particularly *c*ZR and *c*ZROG. This may reflect spatial overlap between *ENT1* expression and *c*Z- and iP-type CK accumulation in the root epidermis and LRC, where these CK types predominate over *t*Z-type CKs^61^. *IPT2* and *IPT9* are broadly expressed, with strong expression in proliferating tissues^62–64^. By contrast, *t*Z-type CK biosynthesis and long-distance transport are more closely associated with the root vasculature^19,61,65^, where *ENT1* expression is limited.

The absence of a strong constitutive phenotype in *ent1* null mutants does not preclude an important transport function. It suggests that ENT1-dependent CK riboside gatekeeping is most relevant in spatially restricted or physiologically dynamic contexts.

*ENT1* expression in epidermal and other outer root tissues positions it at the plant-microbe interface, where microorganisms can alter host CK status by stimulating CK metabolism or producing CKs and CK-like compounds, including riboside forms^66–68^. Beneficial *Trichoderma* and *Bacillus* species influence root architecture and immunity, with CK signalling implicated in microbe-induced resistance^36,69,70^. CK profiling of 22 *Trichoderma* strains showed that their CK pools were dominated by riboside and nucleotide derivatives of *c*Z and iP^36^. Accordingly, *ent1* mutants showed attenuated growth-related and immune responses to pretreatments with beneficial *Trichoderma* and *Bacillus* strains (Fig. 6, Supplementary Fig. 12). Given that *c*ZROG was consistently elevated in *ent1-4* plants, the link between ENT1 and *c*Z-type CK metabolism may be particularly relevant to microbial responses.

Bioassays and receptor studies generally place *c*Z-type CKs below *t*Z-type forms in potency^71,72^, but potency alone does not capture context-dependent activity. *c*Z-type CKs have been linked to growth-limiting conditions, stress responses and biotic interactions^72,73^. In tobacco, *c*Z showed greater early suppression of visible *Pseudomonas syringae* symptoms than *t*Z, whereas *t*Z showed stronger later symptom suppression and more persistent restriction of pathogen proliferation^73^. These observations support the view that *c*Z is not merely a weaker version of *t*Z, but may tune immune-related outputs in a manner compatible with priming rather than full defence activation.

This interpretation fits beneficial plant-microbe interactions, in which host defence must be potentiated without triggering immune activation strong enough to impair colonization or growth. *c*Z-type CKs or *c*Z-associated metabolic states could contribute to this balance.

We therefore propose that microbial exposure reshapes CK riboside pools through changes in host CK metabolism, contributions from microbially derived CKs, or both, with their intracellular partitioning modulated by ENT1 (Fig. 7). Loss of *ENT1* may disrupt this partitioning, favouring the accumulation or O-glucosylation of riboside-derived CKs and thereby attenuating microbial effects on root architecture and immune priming.

**Fig. 7.**
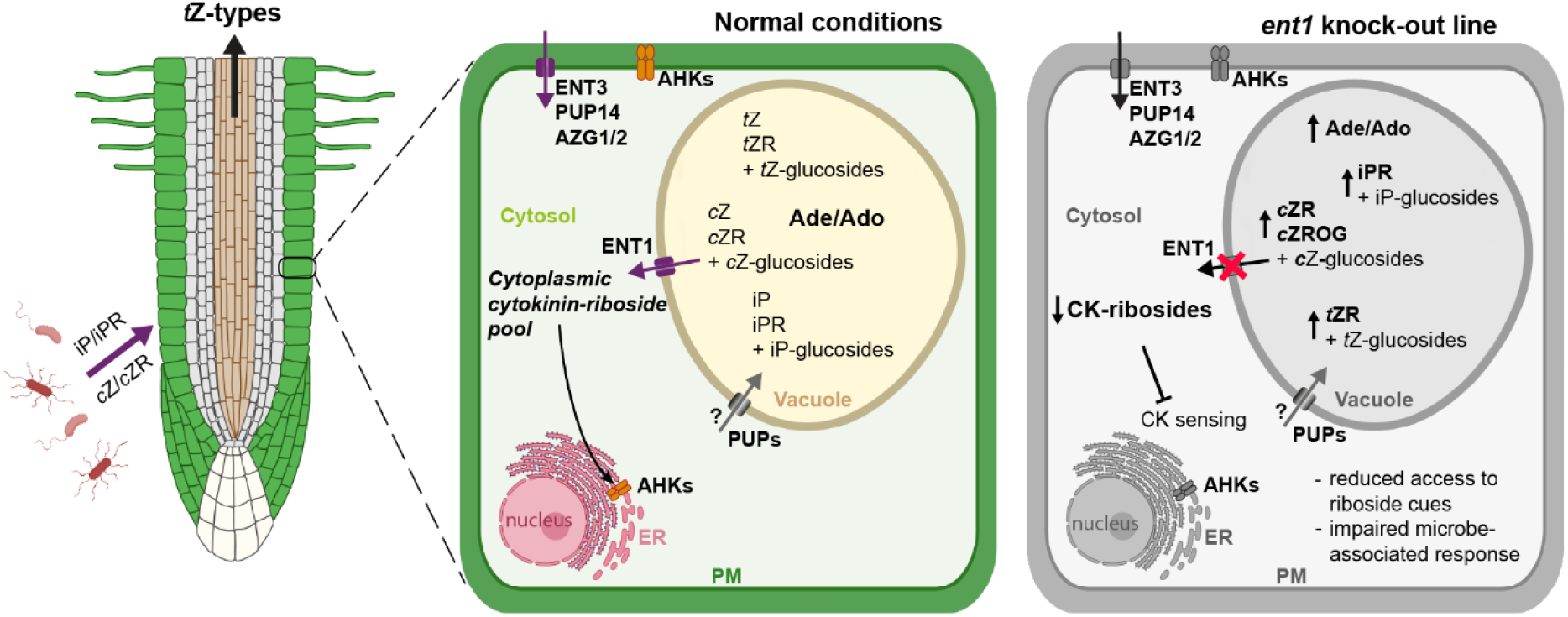
Model for ENT1-dependent cytokinin riboside gatekeeping at the root–microbe interface. ENT1 is enriched in outer root cell layers and localizes predominantly to the tonoplast, where it is proposed to retrieve cytokinin ribosides from the vacuole to the cytosol. This activity may maintain the cytosolic availability of riboside-derived cytokinin pools, particularly *c*Z-type metabolites, and thereby support cytokinin sensing and microbe-associated signalling outputs. In *ent1* mutant plants, impaired vacuolar remobilization is proposed to shift cytokinin riboside pools toward retention or O-glucosylation, reducing their availability for cytosolic cytokinin signalling and thereby affecting beneficial microbe-associated developmental and protective outputs. CK, cytokinin; *t*Z, *trans*-zeatin; *t*ZR, *trans*-zeatin riboside; *c*Z, *cis*-zeatin; *c*ZR, *cis*-zeatin riboside; *c*ZROG, *cis*-zeatin riboside O-glucoside; iP, isopentenyladenine; iPR, isopentenyladenosine; Ade, adenine; Ado, adenosine; ENT, equilibrative nucleoside transporter; PUP, purine permease; AZG, azaguanine resistant; AHK, Arabidopsis histidine kinase; PM, plasma membrane; ER, endoplasmic reticulum.

Together, our results establish ENT1 as a selective tonoplast CK riboside transporter and uncover vacuolar CK riboside retrieval as a regulatory layer in CK biology. By linking intracellular CK partitioning to beneficial microbe-associated protection, this work extends CK transport beyond long-distance movement and receptor-proximal signalling, and identifies vacuolar gatekeeping as a mechanism through which plants tune hormone availability at the plant-microbe interface.

## MATERIAL AND METHODS

### Plant material

Unless stated otherwise, all *Arabidopsis thaliana* (L.) Heynh. lines generated in this study were in the Col-8 background. The *ent1-1* (SALK_025174) and *ent1-2* (SALK_104866) lines were obtained from the Nottingham Arabidopsis Stock Centre (NASC). *ent1-3* and *ent1-4* knockout lines*, pENT1::ENT1^WT^-eGFP:tENT1*, *pENT1::ENT1^L45G^-eGFP:tENT1*, *XVE>>ENT1^WT^*, *XVE>>ENT1^WT^-eGFP*, and *pENT1::NLS-eGFP-GUS:tENT1*-expressing lines were generated as described below.

### Growth conditions

For microscopic and phenotypic analyses, seeds were surface sterilized in 70% ethanol for 3 min, sown on plates containing half-strength Murashige and Skoog (MS) medium (Duchefa; 2.2 g L^−1^ MS salts, 5 mM 2-(N-morpholino)ethanesulfonic acid, pH 5.7), and stratified at 4 °C in the dark for 2 to 3 days.

Seedlings were then grown on vertically oriented plates in a growth chamber (Polyklima) at 21 °C under long-day conditions (16 h light at 120 μmol m^−2^ s^−1^/8 h dark), unless stated otherwise.

For microbial immunity assays, Arabidopsis plants were grown in 7 × 7 cm pots for 3 to 4 weeks at 23 °C and 60% relative humidity under long-day conditions (16 h light at 100 to 120 μmol m^−2^ s^−1^/8 h dark) before microbial interaction experiments.

Tobacco BY-2 (*Nicotiana tabacum* L. cv. Bright Yellow 2) cell lines were maintained in liquid MS medium containing 30 g L^−1^ sucrose, 4.34 g L^−1^ MS salts, 100 mg L^−1^ myo-inositol, 1 mg L^−1^ thiamine, 0.2 mg L^−1^ 2,4-dichlorophenoxyacetic acid and 200 mg L^−1^ KH₂PO₄, adjusted to pH 5.8 and supplemented with 100 mg L^−1^ cefotaxime and 20 mg L^−1^ hygromycin. Cell suspensions were maintained in the dark at 27 °C under continuous shaking on an orbital shaker (150 rpm, orbital diameter 30 mm) and subcultured every 7 days.

### Construct design and molecular cloning

All primer sequences are listed in Supplementary Table 1, where the individual primer pairs are arranged in consecutive rows.

To generate the β-estradiol-inducible *XVE>>ENT1^WT^*construct in *pMDC7*^40^, the *ENT1* genomic sequence was amplified from Arabidopsis genomic DNA using primers ENT1_GWFW01 and ENT1_GWRE01, introduced into *pDONR207* by Gateway BP recombination, and subsequently transferred into *pMDC7* by Gateway LR Clonase II (Thermo Fisher Scientific). The *XVE>>ENT1^WT^-eGFP* construct was generated analogously, except that the *ENT1^WT^-eGFP* fragment was amplified from *pENT1::ENT1^WT^-eGFP pENTR2B* (described below).

Mutant *XVE* constructs carrying the M107Q or Y106S substitutions were derived from the *ENT1^WT^-eGFP* entry clone in *pDONR207* by PCR amplification of overlapping fragments containing the desired mutation, followed by NEBuilder HiFi DNA Assembly (New England Biolabs). For the Y106S:D177S construct, the previously generated Y106S entry clone was used as the template. The resulting entry clones were subsequently recombined into *pMDC7* by Gateway LR Clonase II.

The *pENT1::ENT1^WT^-eGFP:tENT1* construct in *pMCS:GW*^74^ was generated in several consecutive steps. First, the *pENT1::ENT1^WT^-eGFP* construct was assembled in *pENTR2B-GFP*^75^ using a restriction-ligation strategy with *Sal*I and *Not*I. For this, an *ENT1* genomic fragment containing a 4 kb upstream promoter region and the genomic *ENT1* sequence was amplified and cloned into the *pENTR2B-GFP* backbone, yielding a C-terminal *ENT1^WT^-eGFP* fusion in *pENTR2B*. Next, the 840 bp *ENT1* terminator region was added downstream of eGFP by NEBuilder HiFi DNA Assembly to generate C-terminal *pENT1::ENT1^WT^-eGFP:tENT1* in *pENTR2B*.

This construct was then used to generate an untagged *pENT1::ENT1^WT^:tENT1* entry clone by replacing the C-terminal *ENT1^WT^-eGFP* fragment with the genomic *ENT1* sequence amplified from *XVE>>ENT1^WT^ pMDC7*.

Subsequently, eGFP was inserted in frame between codons Gln22 and Met23 by PCR amplification of the eGFP fragment and linearization of the *pENT1::ENT1^WT^:tENT1* backbone at the insertion site, followed by NEBuilder assembly. The *pENT1::ENT1^WT^-eGFP:tENT1* cassette was transferred from *pENTR2B* into *pMCS:GW* by LR recombination.

The *pENT1::ENT1^L45G^-eGFP:tENT1* construct was generated from *pENT1::ENT1^WT^-eGFP:tENT1 pENTR2B* by introducing the L45G substitution through PCR amplification of overlapping fragments followed by NEBuilder assembly. The *pENT1::ENT1^L45G^-eGFP:tENT1* cassette was transferred from *pENTR2B* into *pMCS:GW* by LR recombination.

The *pENT1::NLS-eGFP-GUS:tENT1* construct in *pMCS:GW* was also generated by sequential PCR and NEBuilder assembly. First, the *ENT1* putative terminator was added to the *pENTR2B NLS-eGFP-GUS* backbone^76^. Next, the 4 kb *ENT1* putative promoter region was inserted upstream of the reporter cassette. The *pENT1::NLS-eGFP-GUS:tENT1* cassette was then transferred from *pENTR2B* into *pMCS:GW* by LR recombination.

The CRISPR-Cas9 construct was generated using Golden Gate cloning^77^. The sgRNA cassette was first amplified from *pCBC-DT1T2*^78^, re-amplified to introduce *Bsa*I recognition sites, and finally assembled into *pCF588* by *Bsa*I-mediated Golden Gate cloning.

### Plant transformation

Transgenic and CRISPR/Cas9 lines generated in this study were produced in the *Arabidopsis thaliana* Col-8 background by floral dip^79^ using *Agrobacterium tumefaciens* GV3101 carrying the corresponding construct. Primary transformants (T1) harbouring *pENT1::ENT1^WT^-eGFP:tENT1*, *pENT1::ENT1^L45G^-eGFP:tENT1* or *pENT1::NLS-eGFP-GUS:tENT1* transgenes were selected on soil by spraying with 0.02% Basta (glufosinate-ammonium; Bayer), and subsequent generations were selected on ½ MS medium supplemented with 15 mg L^−1^ phosphinothricin (glufosinate-ammonium, 95%; Thermo Fisher). Transformants carrying the CRISPR/Cas9 construct *pCF588* with ENT1-targeting sgRNAs were selected based on seed eGFP fluorescence. *XVE>>ENT1^WT^* and *XVE>>ENT1^WT^-eGFP* lines were selected on ½ MS medium supplemented with 15 mg L^−1^ hygromycin B (Roche) from the T1 generation onwards.

### Transformation of BY-2 cells

BY-2 cells were transformed with *XVE>>ENT1^WT^-eGFP*, *XVE>>ENT1^Y^*^106^*^S^-eGFP*, *XVE>>ENT1^Y^*^106^*^S:D^*^177^*^S^-eGFP*, and *XVE>>ENT1^M^*^107^*^Q^-eGFP* constructs by co-cultivation with *Agrobacterium tumefaciens* strain GV2260^80^. Transgenic lines were harvested after 4 weeks, cultured on kanamycin-containing solid medium, and tested for the presence of *ENT1* by PCR.

### Hypocotyl explant assay

Plants were cultivated for 1 day in the light and 5 days in the dark on full-strength MS medium with Gamborg B5 vitamins and 10 g L^−1^ sucrose in growth chambers (Polyklima) at 21 °C. Hypocotyls excised from these seedlings were placed on full-strength MS medium with Gamborg B5 vitamins, 10 g L^−1^ sucrose, 1 mg L^−1^ biotin, 300 µg L^−1^ *t*ZR, and 100 µg L^−1^ 1-naphthylacetic acid. Hypocotyl explants were cultivated for 21 days under long-day conditions (16 h light at 21 °C and 8 h dark at 19 °C), at a light intensity of 100 μmol m^−2^ s^−1^. To measure the area of regenerated tissues, Petri dishes were imaged from above using an Epson Perfection V700 Photo scanner (Epson) at a resolution of 800 dots per inch. Image regions corresponding to the regenerated explants were segmented in the hue channel using Otsu thresholding^81^. Binary masks resulting from the segmentation were cleaned by removing regions smaller than 3,000 pixels and larger than 15,000 pixels. Additional segmentation artefacts were removed manually. Properties of the remaining regions were measured. Image processing was performed using the “scikit-image” Python library^82^.

### Plant hormone quantification

Samples for hormone quantification were treated with 1 M formic acid and homogenized using the FastPrep-24 grinder (MP Biomedicals). The homogenate was centrifuged at 30,130 × *g* (17,500 rpm) for 25 min at 4 °C using a 5430R centrifuge (Eppendorf). The supernatant was collected, the pellet was resuspended in 1 M formic acid and centrifugation was repeated as described above. The combined supernatants were loaded onto the Oasis HLB 96-well µElution plates (Waters) that had been previously activated with acetonitrile, ultrapure water, and 1 M formic acid. The plate wells were washed with ultrapure water, and the plant hormone-containing fraction was eluted with 50% acetonitrile, using the Pressure+ manifold (Biotage). The eluted fraction was analysed using a 1290 Infinity II UHPLC system coupled to a 6495 triple quadrupole mass spectrometer (Agilent).

### Uptake and transport of radiolabelled *t*ZR

Arabidopsis plants were grown on ½ MS medium without sucrose. Eight-day-old plants were placed into the wells of 24-well plates (three plants per well in eight replicates) filled with 0.90 mL of [2-³H]*t*ZR solution so that the root tips were submerged, but the root bases and aerial tissues were not. Plants were treated for 4 h, during which the plates were covered and sealed with Parafilm. After treatment, shoots and roots were separated, placed individually into glass scintillation vials, and 100 µL of 5% sodium hypochlorite (NaOCl) solution was added. After 16 h, the samples were thoroughly mixed with 5 mL of scintillation cocktail (Rotiszint Flowcount), and liquid scintillation counting was performed on a Tri-Carb 4910 TR (PerkinElmer). [2-³H]*t*ZR was synthesized in the Isotope Laboratory of the Institute of Experimental Botany of the Czech Academy of Sciences. The purity of the stock solution used in the experiment was over 95%, as determined by thin-layer chromatography. After dilution with distilled water, the solution used in the experiment had an activity of 79.21 kBq mL^−1^ and a concentration of 64.81 nM.

### Imaging

Imaging was performed using a Zeiss LSM900 microscope equipped with Plan-Apochromat 10×/0.45 M27, Plan-Apochromat 20×/0.8 M27 and LD LCI Plan-Apochromat 40×/1.2 Imm Korr DIC M27 objectives. Detection was performed using standard GaAsP-PMTs and the multi-channel Airyscan 2 detector (Zeiss). Images were captured using excitation wavelengths of 488 nm for eGFP, 561 nm for FM4-64 and propidium iodide, and 640 nm for LysoTracker Deep Red and chlorophyll autofluorescence. ZEN Blue software (Zeiss) and Fiji were used for image post-processing.

### Protein extraction, SDS-PAGE, western blotting and immunodetection

Roots from 14-day-old seedlings were homogenized in lysis buffer (100 mM Tris-HCl, 5% SDS, 100 mM NaCl, 1 mM DTT, pH 8.0) at a 1:1 (w/v) ratio. The lysis buffer was supplemented with cOmplete protease inhibitor cocktail (Roche) to prevent protein degradation. The crude extract was centrifuged twice at 10,000 × *g* for 10 min at 4 °C, and the cleared protein extract was collected for subsequent analysis. Proteins were separated on 10% SDS-PAGE gels (Bio-Rad) under denaturing conditions and transferred to a PVDF membrane (Millipore) using a wet transfer system. Transfer efficiency was verified by reversible staining with Ponceau S.

For immunodetection, membranes were probed with the following primary antibodies: anti-ENT1, a custom polyclonal affinity-purified rabbit antibody (Moravian-Biotech) raised against the peptide sequence FHEERKNEELIREKS of the ENT1 protein, used at 1:7,000 dilution; and anti-actin, a commercial plant-specific mouse monoclonal antibody (ABclonal), used at 1:5,000 dilution.

Primary antibodies were diluted in 5% non-fat milk in TTBS, which was also used for blocking and diluting the secondary antibody. Horseradish peroxidase (HRP)-conjugated goat anti-mouse IgG (H+L) and goat anti-rabbit IgG (H+L) secondary antibodies (ABclonal) were applied at a 1:5,000 dilution. Immunoreactive bands were visualized using enhanced chemiluminescence (ECL; Amersham).

### BY-2 radioaccumulation assay

Radioaccumulation assays and mathematical modelling of cellular uptake rate constants were performed as described previously^28^. *ENT1*-transformed BY-2 cell suspensions were treated with either 1 µM β-estradiol in dimethyl sulfoxide (DMSO) or the solvent alone to obtain induced and non-induced samples, respectively. The data points were fitted with the equation:

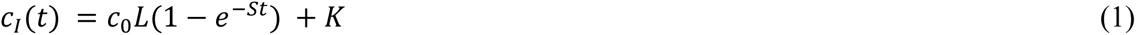

where *c_I_* is the intracellular concentration of the radiolabelled tracer, *t* is time, *c_0_* is the initial tracer concentration in the medium, and *L*, *S*, and *K* are optimizable parameters. The first-order rate constant for tracer uptake, *I*, was calculated as:

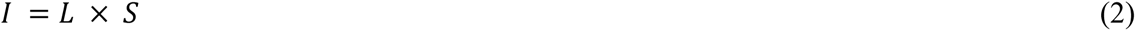

### Molecular docking

We used the AutoDockFR (ADFR) software suite^83,84^ to dock *t*Z into AlphaFold-predicted structural models^85^ of ENT1 and ENT3. The ligand was initially placed in the central cavity of each protein. The centre and dimensions of the affinity grids were determined automatically by the ADFR program “agfr” in ligand mode, with a padding of 0.8 nm. For each docking, 100 runs were performed, with a maximum of 50 million evaluations per run. Mutations were simulated by amino acid replacement in UCSF ChimeraX^86^. Ligand-interacting residues Met107/Gln107 and Gln181 in ENT1, and Gln62/Met62 and Gln133 in ENT3, were defined as flexible.

### RT-qPCR

Total RNA was extracted from 10-day-old *Arabidopsis thaliana* seedlings using the Spectrum Plant Total RNA Kit (Sigma-Aldrich). Residual genomic DNA was removed using the DNA-free DNA Removal Kit (Thermo Fisher Scientific). First-strand cDNA was synthesized from 2 μg of total RNA using RevertAid H Minus Reverse Transcriptase (Thermo Scientific) and an oligo(dT) primer.

The resulting cDNA was diluted and used as a template for reverse transcription quantitative PCR (RT-qPCR). qPCR was performed in 96-well plates on a StepOnePlus Real-Time PCR System (Applied Biosystems) using gb SG PCR Master Mix (Generi Biotech), according to the manufacturers’ instructions. Transcript levels were normalized to selected reference genes, and relative expression was calculated using the ΔΔ*C_t_* method^87^.

### Microbial treatments

*Trichoderma harzianum* isolate T39 was cultured on potato dextrose agar (PDA, Difco lab) plates at 26 ± 2 °C in natural light. T39 conidia were harvested in distilled water and adjusted to a concentration of 10^7^ conidia mL^−1^. 2 mL of the conidia suspension was applied to plants *via* soil drench^70^.

*Bacillus megaterium* strain 4C and *Bacillus pumilus* strain R2E^35^ were grown overnight in Luria-Bertani broth (LB; Difco lab) at 26 ± 2 °C on a shaker at 120 rpm. After incubation, bacterial cultures were individually centrifuged at 12,000 rpm for 5–8 min. The bacterial cultures were washed twice in distilled water, and re-suspended in sterile distilled water. The cell suspension was adjusted to an optical density of OD_600_ = 0.1 (∼10^8^ CFU mL^−1^), and 2 mL of bacterial suspension was applied to plants via soil drench^35^.

For pathogenesis and immunity assays, T39, 4C, and R2E inocula were applied to 4- to 5-week-old plants by soil drench per pot at the indicated concentrations. Two treatments were applied: 4 days and 24 h before the pathogen challenge.

### Plant infection

*Botrytis cinerea* strain B05.10 was subcultured on PDA plates and incubated at 22 ± 2 °C for 3–4 days. For infection, agar discs measuring 0.4 cm in diameter were cut from the colony margins using a cork borer and placed on detached leaves. The leaves were kept in a humid chamber at 22 ± 1 °C and maintained under long-day conditions. *B. cinerea* lesion size was assessed 3 days post-inoculation using ImageJ software^88^.

*Pseudomonas syringae* pv. tomato DC3000 was grown in LB broth containing 100 mg L^−1^ rifampicin overnight at 28 °C. Overnight log phase cultures were pelleted, washed three times with 10 mM MgCl_2_, standardized to OD_600_ = 0.2, and diluted to a final OD_600_ = 0.0002 for inoculation by syringe infiltration. Mature second- and third-tier rosette leaves from 4-week-old plants were infiltrated with the bacterial suspension. Three leaves per plant were infiltrated on the abaxial side with the bacterial suspension using a 1 mL blunt-tipped syringe. Mock-treated plants were similarly inoculated with 10 mM MgCl_2_ in the absence of bacteria. Three days following infiltration, three leaf discs of 0.6 cm diameter were excised from at least four plants from each genotype, and macerated in 1 mL of 10 mM MgCl_2_ to release intercellular bacteria. For plating of serial dilutions (from 10^−1^ to 10^−6^), 100 μL of each dilution was spread on an LB plate containing 100 mg L^−1^ rifampicin. The plates were incubated at 28 °C for 40 h, and the resulting colonies were counted to determine bacterial colony-forming units (CFU)^89,90^. Disease was also assessed by quantifying the water-soaked lesion area in ImageJ.

### Reactive oxygen species measurement

Reactive oxygen species (ROS) generation was measured as previously described^91,92^. Leaf discs 0.5 cm in diameter were harvested from Arabidopsis plants treated with the microbial inocula described above. Leaf discs were floated in wells of a white 96-well plate (SPL Life Sciences) containing 300 μL of distilled water at room temperature. After 24 h of incubation, the water was removed, and 50 μL of reaction mixture containing 150 μM luminol, 15 μg mL^−1^ HRP, and 1 μM flg22 peptide was immediately added. Light emission was measured immediately using a GloMax microplate luminometer (Promega).

### Statistical analysis

Statistical analyses were performed using the tests specified in the corresponding figure legends. Pairwise comparisons of independent samples were analysed using Welch’s two-sample t-test or Student’s two-tailed t-test, as appropriate. Paired datasets were analysed using a paired t-test. Non-parametric pairwise comparisons were analysed using a two-tailed Mann-Whitney U test. Multiple-group comparisons were analysed using the Kruskal-Wallis test, the Kruskal-Wallis test followed by Dunn’s post hoc test, or Welch’s ANOVA followed by Dunnett’s post hoc test, as appropriate. Data are presented as individual data points with boxplots showing the median and interquartile range (IQR), unless otherwise stated. Whiskers indicate either the most extreme non-outlier values within 1.5 × IQR or the full data range, as specified in the corresponding figure legends. For qRT-PCR analyses, data are shown as Δ*C_t_* values or mean values as specified in the corresponding figure legends. Differences were considered statistically significant at *P* < 0.05.

## ACKNOWLEDGMENTS

The work was supported by the Czech Science Foundation via project 26-23352S (M.H., D.N., A.K., P.K., M.K., K.H., and O.P.), European Regional Development Fund-Project CZ.02.01.01/00/23_020/0008497 (SMART Plant Biotechnology for Sustainable Agriculture, O.P.), and by ERDF project CZ.02.02.01/00/23_023/0009111 (Development of educational infrastructure and innovative approaches to teaching at Palacký University in Olomouc, J.Š. and O.N.). HPLC/MS analysis of plant hormones was performed at the Bioanalytical Service Laboratory of the IEB AS CR. Computational resources were provided by the e-INFRA CZ project (ID:90254), supported by the Ministry of Education, Youth and Sports of the Czech Republic. BY2-related microscopic observations were performed at the Imaging Facility of the IEB AS CR, supported by MEYS CR LM2023050 ‘Czech-BioImaging’ and IEB AS CR. We thank Lenka Drašarová for the synthesis of [2-³H]*t*ZR and Dekel Cohen-Hoch for helpful technical advice and training.

## AUTHOR CONTRIBUTIONS

M.H., O.P., D.N., and K.H. conceptualized and designed the study. M.H., A.K., V.S., and L.T.P. performed molecular cloning and most of the Arabidopsis work, including microscopic techniques. D.N., M.K., and K.H. performed all BY-2 transport experiments, and D.N. performed in silico analyses. D.N., P.K., M.K., and S.T.F. conducted radioactivity and hypocotyl assays. R.G., L.T., and M.B. performed all microbe treatments and ROS measurements. T.M. and D.Z. performed genotyping and expression analysis of *ent1* mutants. J.Š., O.Š., and J.H. provided part of the infrastructure and expert consultations with microscopy and plant cultivation. O.N., J.Š., O.Š., T.M., and E.B. contributed to the formal analysis. M.H., D.N., and O.P. wrote the manuscript with input from all co-authors. All authors read and edited the manuscript.

## DATA AVAILABILITY STATEMENT

All data supporting the findings of this study are available within the article and its Supplementary Information files. Source data are provided with this paper. Additional raw data are available from the corresponding authors upon reasonable request.

## CODE AVAILABILITY STATEMENT

Custom scripts used for image segmentation, curve fitting, and downstream quantitative analyses are available from the corresponding authors upon reasonable request.

## REFERENCES

1. Kieber, J. J. & Schaller, G. E. Cytokinin signaling in plant development. Development 145, dev149344 (2018).

2. Cortleven, A. & Schmülling, T. Regulation of chloroplast development and function by cytokinin. J. Exp. Bot. 66, 4999–5013 (2015).

3. Kábrtová, V. et al. Cytokinin downregulates Photosystem II photochemistry during prolonged darkness in a phytochrome B-dependent manner. New Phytol. 250, 3846–3866 (2026).

4. Großkinsky, D. K. et al. Cytokinins mediate resistance against *Pseudomonas syringae* in tobacco through increased antimicrobial phytoalexin synthesis independent of salicylic acid signaling. Plant Physiol. 157, 815–830 (2011).

5. Lomin, S. N. et al. Plant membrane assays with cytokinin receptors underpin the unique role of free cytokinin bases as biologically active ligands. J. Exp. Bot. 66, 1851–1863 (2015).

6. Suzuki, T. et al. The *Arabidopsis* sensor His-kinase, AHK4, can respond to cytokinins. Plant Cell Physiol. 42, 107–113 (2001).

7. Inoue, T. et al. Identification of CRE1 as a cytokinin receptor from *Arabidopsis*. Nature 409, 1060–1063 (2001).

8. Romanov, G. A., Lomin, S. N. & Schmülling, T. Biochemical characteristics and ligand-binding properties of *Arabidopsis* cytokinin receptor AHK3 compared to CRE1/AHK4 as revealed by a direct binding assay. J. Exp. Bot. 57, 4051–4058 (2006).

9. Kurakawa, T. et al. Direct control of shoot meristem activity by a cytokinin-activating enzyme. Nature 445, 652–655 (2007).

10. Kuroha, T. et al. Functional analyses of LONELY GUY cytokinin-activating enzymes reveal the importance of the direct activation pathway in *Arabidopsis*. Plant Cell 21, 3152–3169 (2009).

11. Werner, T. et al. Cytokinin-deficient transgenic *Arabidopsis* plants show multiple developmental alterations indicating opposite functions of cytokinins in the regulation of shoot and root meristem activity. Plant Cell 15, 2532–2550 (2003).

12. Hou, B., Lim, E.-K., Higgins, G. S. & Bowles, D. J. N-glucosylation of cytokinins by glycosyltransferases of *Arabidopsis thaliana*. J. Biol. Chem. 279, 47822–47832 (2004).

13. Wang, J., Ma, X.-M., Kojima, M., Sakakibara, H. & Hou, B.-K. N-glucosyltransferase UGT76C2 is involved in cytokinin homeostasis and cytokinin response in *Arabidopsis thaliana*. Plant Cell Physiol. 52, 2200–2213 (2011).

14. Šmehilová, M., Dobrůšková, J., Novák, O., Takáč, T. & Galuszka, P. Cytokinin-specific glycosyltransferases possess different roles in cytokinin homeostasis maintenance. Front. Plant Sci. 7, 1264 (2016).

15. Brzobohatý, B. et al. Release of active cytokinin by a β-glucosidase localized to the maize root meristem. Science 262, 1051–1054 (1993).

16. Osugi, A. et al. Systemic transport of trans-zeatin and its precursor have differing roles in *Arabidopsis* shoots. Nat. Plants 3, 17112 (2017).

17. Landrein, B. et al. Nitrate modulates stem cell dynamics in *Arabidopsis* shoot meristems through cytokinins. Proc. Natl Acad. Sci. USA 115, 1382–1387 (2018).

18. Zhang, K., et al. *Arabidopsis* ABCG14 protein controls the acropetal translocation of root-synthesized cytokinins. Nat. Commun. 5, 3274 (2014).

19. Ko, D., et al. *Arabidopsis* ABCG14 is essential for the root-to-shoot translocation of cytokinin. Proc. Natl Acad. Sci. USA 111, 7150–7155 (2014).

20. Tessi, T. M., et al. *Arabidopsis* AZG2 transports cytokinins in vivo and regulates lateral root emergence. New Phytol. 229, 979–993 (2021).

21. Tessi, T. M. et al. AZG1 is a cytokinin transporter that interacts with auxin transporter PIN1 and regulates the root stress response. New Phytol. 238, 1924–1941 (2023).

22. Li, G., Liu, K., Baldwin, S. A. & Wang, D. Equilibrative nucleoside transporters of *Arabidopsis thaliana*. cDNA cloning, expression pattern, and analysis of transport activities. J. Biol. Chem. 278, 35732–35742 (2003).

23. Wormit, A., Traub, M., Flörchinger, M., Neuhaus, H. E. & Möhlmann, T. Characterization of three novel members of the *Arabidopsis thaliana* equilibrative nucleoside transporter (ENT) family. Biochem. J. 383, 19–26 (2004).

24. Girke, C., Daumann, M., Niopek-Witz, S. & Möhlmann, T. Nucleobase and nucleoside transport and integration into plant metabolism. Front. Plant Sci. 5, 443 (2014).

25. Sun, J., et al. *Arabidopsis* SOI33/AtENT8 gene encodes a putative equilibrative nucleoside transporter that is involved in cytokinin transport in planta. J. Integr. Plant Biol. 47, 588–603 (2005).

26. Hirose, N. et al. Regulation of cytokinin biosynthesis, compartmentalization and translocation. J. Exp. Bot. 59, 75–83 (2008).

27. Korobova, A., Kuluev, B., Möhlmann, T., Veselov, D. & Kudoyarova, G. Limitation of cytokinin export to the shoots by nucleoside transporter ENT3 and its linkage with root elongation in *Arabidopsis*. Cells 10, 350 (2021).

28. Nedvěd, D. et al. Membrane transport of root-borne trans-zeatin riboside maintains the cytokinin homeostasis in shoots. J. Exp. Bot. 76, 6723–6740 (2025).

29. Möhlmann, T., Mezher, Z., Schwerdtfeger, G. & Neuhaus, H. E. Characterisation of a concentrative type of adenosine transporter from *Arabidopsis thaliana* (ENT1,At). FEBS Lett. 509, 370–374 (2001).

30. Bernard, C. et al. Equilibrative nucleoside transporter 1 (ENT1) is critical for pollen germination and vegetative growth in *Arabidopsis*. J. Exp. Bot. 62, 4627–4637 (2011).

31. Kowalska, M. et al. Vacuolar and cytosolic cytokinin dehydrogenases of *Arabidopsis thaliana*: Heterologous expression, purification and properties. Phytochemistry 71, 1970–1978 (2010).

32. Kiran, N. S. et al. Retargeting a maize β-glucosidase to the vacuole: evidence from intact plants that zeatin-*O*-glucoside is stored in the vacuole. Phytochemistry 79, 67–77 (2012).

33. Jiskrová, E. et al. Extra- and intracellular distribution of cytokinins in the leaves of monocots and dicots. New Biotechnol. 33, 735–742 (2016).

34. Skalický, V., Kubeš, M., Napier, R. & Novák, O. Auxins and cytokinins: the role of subcellular organization on homeostasis. Int. J. Mol. Sci. 19, 3115 (2018).

35. Gupta, R. et al. Cytokinin drives assembly of the phyllosphere microbiome and promotes disease resistance through structural and chemical cues. ISME J. 16, 122–137 (2022).

36. Bean, K. M., Kisiala, A. B., Morrison, E. N. & Emery, R. J. N. *Trichoderma* synthesizes cytokinins and alters cytokinin dynamics of inoculated *Arabidopsis* seedlings. J. Plant Growth Regul. 41, 2678–2694 (2022).

37. Gupta, R., Elkabetz, D., Leibman-Markus, M., Jami, E. & Bar, M. Cytokinin-microbiome interactions regulate developmental functions. *Environ*. Microbiome 17, 2 (2022).

38. Großkinsky, D. K. et al. Cytokinin production by *Pseudomonas fluorescens* G20-18 determines biocontrol activity against *Pseudomonas syringae* in *Arabidopsis*. Sci. Rep. 6, 23310 (2016).

39. Zuo, J., Niu, Q. W. & Chua, N. H. Technical advance: An estrogen receptor-based transactivator XVE mediates highly inducible gene expression in transgenic plants. Plant J. 24, 265–273 (2000).

40. Curtis, M. D. & Grossniklaus, U. A gateway cloning vector set for high-throughput functional analysis of genes in planta. Plant Physiol. 133, 462–469 (2003).

41. Li, J. & Wang, D. Cloning and in vitro expression of the cDNA encoding a putative nucleoside transporter from *Arabidopsis thaliana*. Plant Sci. 157, 23–32 (2000).

42. Jaquinod, M. et al. A proteomics dissection of *Arabidopsis thaliana* vacuoles isolated from cell culture. Mol. Cell. Proteomics 6, 394–412 (2007).

43. Wolfenstetter, S., Wirsching, P., Dotzauer, D., Schneider, S. & Sauer, N. Routes to the tonoplast: the sorting of tonoplast transporters in *Arabidopsis* mesophyll protoplasts. Plant Cell 24, 215–232 (2012).

44. Wang, X. et al. Trans-Golgi network-located AP1 gamma adaptins mediate dileucine motif-directed vacuolar targeting in *Arabidopsis*. Plant Cell 26, 4102–4118 (2014).

45. Kudo, T., Kiba, T. & Sakakibara, H. Metabolism and long-distance translocation of cytokinins. J. Integr. Plant Biol. 52, 53–60 (2010).

46. Kuderová, A. et al. Effects of conditional IPT-dependent cytokinin overproduction on root architecture of *Arabidopsis* seedlings. Plant Cell Physiol. 49, 570–582 (2008).

47. Zubo, Y. O., et al. Cytokinin induces genome-wide binding of the type-B response regulator ARR10 to regulate growth and development in *Arabidopsis*. Proc. Natl Acad. Sci. USA 114, E5995–E6004 (2017).

48. Steiner, E. et al. Characterization of the cytokinin sensor TCSv2 in *Arabidopsis* and tomato. Plant Methods 16, 152 (2020).

49. Kubo, M. & Kakimoto, T. The CYTOKININ-HYPERSENSITIVE genes of *Arabidopsis* negatively regulate the cytokinin-signaling pathway for cell division and chloroplast development. Plant J. 23, 385–394 (2000).

50. Pernisová, M. et al. Cytokinins modulate auxin-induced organogenesis in plants via regulation of the auxin efflux. Proc. Natl Acad. Sci. USA 106, 3609–3614 (2009).

51. Pelagio-Flores, R., Esparza-Reynoso, S., Garnica-Vergara, A., López-Bucio, J. & Herrera-Estrella, A. *Trichoderma*-induced acidification is an early trigger for changes in *Arabidopsis* root growth and determines fungal phytostimulation. Front. Plant Sci. 8, 822 (2017).

52. Contreras-Cornejo, H. A., Macías-Rodríguez, L., Alfaro-Cuevas, R. & López-Bucio, J. *Trichoderma* spp. improve growth of *Arabidopsis* seedlings under salt stress through enhanced root development, osmolite production, and Na+ elimination through root exudates. Mol. Plant Microbe Interact. 27, 503–514 (2014).

53. Argueso, C. T. et al. Two-component elements mediate interactions between cytokinin and salicylic acid in plant immunity. PLoS Genet. 8, e1002448 (2012).

54. Li, B., Wang, R., Wang, S., Zhang, J. & Chang, L. Diversified regulation of cytokinin levels and signaling during *Botrytis cinerea* infection in *Arabidopsis*. Front. Plant Sci. 12, 584042 (2021).

55. Paasch, B. C. et al. A critical role of a eubiotic microbiota in gating proper immunocompetence in *Arabidopsis*. Nat. Plants 9, 1468–1480 (2023).

56. Arnaud, D. et al. Cytokinin-mediated regulation of reactive oxygen species homeostasis modulates stomatal immunity in *Arabidopsis*. Plant Cell 29, 543–559 (2017).

57. Kubiasová, K., et al. Cytokinin fluoroprobe reveals multiple sites of cytokinin perception at plasma membrane and endoplasmic reticulum. Nat. Commun. 11, 4285 (2020).

58. Antoniadi, I. et al. Cell-surface receptors enable perception of extracellular cytokinins. Nat. Commun. 11, 4284 (2020).

59. Wulfetange, K. et al. The cytokinin receptors of *Arabidopsis* are located mainly to the endoplasmic reticulum. Plant Physiol. 156, 1808–1818 (2011).

60. Caesar, K. et al. Evidence for the localization of the *Arabidopsis* cytokinin receptors AHK3 and AHK4 in the endoplasmic reticulum. J. Exp. Bot. 62, 5571–5580 (2011).

61. Antoniadi, I. et al. Cell-type-specific cytokinin distribution within the *Arabidopsis* primary root apex. Plant Cell 27, 1955–1967 (2015).

62. Miyawaki, K., Matsumoto-Kitano, M. & Kakimoto, T. Expression of cytokinin biosynthetic isopentenyltransferase genes in *Arabidopsis*: tissue specificity and regulation by auxin, cytokinin, and nitrate. Plant J. 37, 128–138 (2004).

63. Miyawaki, K. et al. Roles of *Arabidopsis* ATP/ADP isopentenyltransferases and tRNA isopentenyltransferases in cytokinin biosynthesis. Proc. Natl Acad. Sci. USA 103, 16598–16603 (2006).

64. Antoniadi, I. et al. IPT9, a cis-zeatin cytokinin biosynthesis gene, promotes root growth. Front. Plant Sci. 13, 932008 (2022).

65. Kiba, T., Takei, K., Kojima, M. & Sakakibara, H. Side-chain modification of cytokinins controls shoot growth in *Arabidopsis*. Dev. Cell 27, 452–461 (2013).

66. Akhtar, S. S., Mekureyaw, M. F., Pandey, C. & Roitsch, T. Role of cytokinins for interactions of plants with microbial pathogens and pest insects. Front. Plant Sci. 10, 1777 (2020).

67. García de Salamone, I. E., Hynes, R. K. & Nelson, L. M. Cytokinin production by plant growth promoting rhizobacteria and selected mutants. Can. J. Microbiol. 47, 404–411 (2001).

68. Illescas, M., Pedrero-Méndez, A., Pitorini-Bovolini, M., Hermosa, R. & Monte, E. Phytohormone production profiles in *Trichoderma* species and their relationship to wheat plant responses to water stress. Pathogens 10, 991 (2021).

69. Poveda, J. & González-Andrés, F. *Bacillus* as a source of phytohormones for use in agriculture. Appl. Microbiol. Biotechnol. 105, 8629–8645 (2021).

70. Leibman-Markus, M. et al. Immunity priming uncouples the growth–defense trade-off in tomato. Development 150, dev201158 (2023).

71. Spíchal, L. et al. Two cytokinin receptors of *Arabidopsis thaliana*, CRE1/AHK4 and AHK3, differ in their ligand specificity in a bacterial assay. Plant Cell Physiol. 45, 1299–1305 (2004).

72. Schäfer, M. et al. The role of *cis*-zeatin-type cytokinins in plant growth regulation and mediating responses to environmental interactions. J. Exp. Bot. 66, 4873–4884 (2015).

73. Großkinsky, D. K., et al. Structure–function relation of cytokinins determines their differential efficiency in mediating tobacco resistance against *Pseudomonas syringae*. Physiol. Plant. 177, e70028 (2025).

74. Michniewicz, M., Frick, E. M. & Strader, L. C. Gateway-compatible tissue-specific vectors for plant transformation. BMC Res. Notes 8, 63 (2015).

75. Králová, M. et al. A decoy receptor derived from alternative splicing fine-tunes cytokinin signaling in *Arabidopsis*. Mol. Plant 17, 1850–1865 (2024).

76. Hajný, J. et al. Sucrose-responsive osmoregulation of plant cell size by a long non-coding RNA. Mol. Plant 17, 1719–1732 (2024).

77. Gao, X. et al. Modular construction of plasmids by parallel assembly of linear vector components. Anal. Biochem. 437, 172–177 (2013).

78. Xing, H.-L. et al. A CRISPR/Cas9 toolkit for multiplex genome editing in plants. BMC Plant Biol. 14, 327 (2014).

79. Clough, S. J. & Bent, A. F. Floral dip: a simplified method for *Agrobacterium*-mediated transformation of *Arabidopsis thaliana*. Plant J. 16, 735–743 (1998).

80. Deblaere, R. et al. Efficient octopine Ti plasmid-derived vectors for *Agrobacterium*-mediated gene transfer to plants. Nucleic Acids Res. 13, 4777–4788 (1985).

81. Liao, P.-S., Chen, T.-S. & Chung, P.-C. A fast algorithm for multilevel thresholding. J. Inf. Sci. Eng. 17, 713–727 (2001).

82. van der Walt, S. et al. scikit-image: image processing in Python. PeerJ 2, e453 (2014).

83. Zhao, Y., Stoffler, D. & Sanner, M. Hierarchical and multi-resolution representation of protein flexibility. Bioinformatics 22, 2768–2774 (2006).

84. Ravindranath, P. A., Forli, S., Goodsell, D. S., Olson, A. J. & Sanner, M. F. AutoDockFR: advances in protein-ligand docking with explicitly specified binding site flexibility. PLoS Comput. Biol. 11, e1004586 (2015).

85. Jumper, J. et al. Highly accurate protein structure prediction with AlphaFold. Nature 596, 583–589 (2021).

86. Goddard, T. D. et al. UCSF ChimeraX: meeting modern challenges in visualization and analysis. Protein Sci. 27, 14–25 (2018).

87. Livak, K. J. & Schmittgen, T. D. Analysis of relative gene expression data using real-time quantitative PCR and the 2^−ΔΔCT^ method. Methods 25, 402–408 (2001).

88. Gupta, R., et al. *Bacillus thuringiensis* promotes systemic immunity in tomato, controlling pests and pathogens and promoting yield. Food Secur. 16, 675–690 (2024).

89. Katagiri, F., Thilmony, R. & He, S. Y. The *Arabidopsis thaliana*-*Pseudomonas syringae* interaction. Arabidopsis Book 1, e0039 (2002).

90. Liu, X. et al. Bacterial leaf infiltration assay for fine characterization of plant defense responses using the *Arabidopsis thaliana*-*Pseudomonas syringae* pathosystem. J. Vis. Exp. e53364 (2015).

91. Leibman-Markus, M., Schuster, S. & Avni, A. LeEIX2 interactors’ analysis and EIX-mediated responses measurement. Methods Mol. Biol. 1578, 167–172 (2017).

92. Anand, G., Leibman-Markus, M., Elkabetz, D. & Bar, M. Method for the production and purification of plant immuno-active xylanase from *Trichoderma*. Int. J. Mol. Sci. 22, 4214 (2021).

