## Supplementary Files for "A tonoplast cytokinin riboside transporter gates intracellular hormone availability at the root–microbe interface"

### SUPPLEMENTARY DATA

#### Supplementary Table 1 | Primer sequences used for molecular cloning, genotyping, and RT-qPCR.

| | |
|---|---|
| **Cloning primers for preparation of *XVE>>ENT1*$^{WT}$ and *XVE>>ENT1*$^{WT}$*-eGFP pMDC7* constructs** | |
| ENT1_GWFW01 | GGGGACAAGTTTGTACAAAAAAGCAGGCTTCATGACTCCCATTAGTCAACGAATTTC |
| ENT1_GWRE01 | GGGGACCACTTTGTACAAGAAAGCTGGGTCTCAAATGACCCAGAACCAAGCAATG |
| **Cloning primers for introduction of the M107Q substitution** | |
| GA_ENT1_FW02 | CTGGCGCCGGAACCAATTCAATGACTCCCATTAGTCAACGAATTTCC |
| GA_ENT1_M107Q_RE01 | CAAACAAGAGCAACGAGCTGGTAGATAACGGC |
| GA_ENT1_M107Q_FW01 | GCCGTTATCTACCAGCTCGTTGCTCTTGTTTG |
| GA_pDONR207_RE02 | CGTTGACTAATGGGAGTCATTGAATTGGTTCCGGCGCC |
| **Cloning primers for introduction of the Y106S substitution** | |
| GA_ENT1_FW02 | CTGGCGCCGGAACCAATTCAATGACTCCCATTAGTCAACGAATTTCC |
| GA_ENT1_Y106S_RE01 | CAAACAAGAGCAACGAGCATGGAGATAACGGCGAAGATCCGAT |
| GA_ENT1_Y106S_FW01 | ATCGGATCTTCGCCGTTATCTCCATGCTCGTTGCTCTTGTTTG |
| GA_pDONR207_RE02 | CGTTGACTAATGGGAGTCATTGAATTGGTTCCGGCGCC |
| **Cloning primers for introduction of the Y106S:D177S substitutions** | |
| GA_ENT1_FW02 | CTGGCGCCGGAACCAATTCAATGACTCCCATTAGTCAACGAATTTCC |
| GA_ENT1_D177S_RE01 | GACCTCCTTGCATAAGTGCAGAACCTAGACCGGATAAGGCAAC |
| GA_ENT1_D177S_FW01 | GTTGCCTTATCCGGTCTAGGTTCTGCACTTATGCAAGGAGGTC |
| GA_pDONR207_RE02 | CGTTGACTAATGGGAGTCATTGAATTGGTTCCGGCGCC |
| **Cloning primers for generation of C-terminal *pENT1::ENT1*$^{WT}$*-eGFP* in *pENTR2B*** | |
| ENT1_prom_SalI_FW04 | GTGTGTGTCGACATGAACATATCAACTCCATAAGTGC |
| ENT1_NotI_RE | GTGTGTGCGGCCGCCCGTCAATGACCCAGAACCAAGCAATG |
| **Cloning primers for introduction of *tENT1* into *pENT1::ENT1*$^{WT}$*-eGFP* in *pENTR2B*** | |
| GA_pENTR2b_ENT1_4kb-GFP_3UTR_FW | TCCTGTCCCAACAAAACCACACTCGAGATATCTAGACCCAGCT |
| GA_pENTR2b_ENT1_4kb-GFP_3UTR_RE | ATAACAAAGCTTTGGTAGAATTACTTGTACAGCTCGTCCATGC |
| GA_ENT1_3UTR_FW | TGGACGAGCTGTACAAGTAATTCTACCAAAGCTTTGTTATCTGTATTTGTTGAA |
| GA_ENT1_3UTR_RE | TGGGTCTAGATATCTCGAGTGTGGTTTTGTTGGGACAGGACG |
| **Cloning primers for generation of untagged *pENT1::ENT1*$^{WT}$*:tENT1* in *pENTR2B*** | |
| GA_pENTR2b_prom_3UTR_FW | CTTGGTTCTGGGTCATTTGATTCTACCAAAGCTTTGTTATCTGTATTTGTTGAA |
| GA_pENTR2b_prom_3UTR_RE | CGTTGACTAATGGGAGTCATTAAAACGGTGCG |
| GA_ENT1_gene_FW | TACAAAACCGCACCGTTTTAATGACTCCCATTAGTCAACGAATTTCC |
| GA_ENT1_gene_RE | ATAACAAAGCTTTGGTAGAATCAAATGACCCAGAACCAAGCAATGACAGAT |
| **Cloning primers for insertion of eGFP between Gln22 and Met23 in *pENT1::ENT1*$^{WT}$*:tENT1*** | |
| GA_ENT1_22_23_GFP_FW | CAAATTCAAATGGTGAGCAAGGGC |
| GA_ENT1_22_23_GFP_RE | GGTGGTCATCTTGTACAGCTCGTCCATG |
| GA_ENT1_22_23_backbone_FW | CTGTACAAGATGACCACCACCGATAAAT |
| GA_ENT1_22_23_backbone_RE | TGCTCACCATTTGAATTTGTTTAGTTTCTCTCAG |
| **Cloning primers for introduction of the L45G substitution in *pENT1::ENT1*$^{WT}$*-eGFP:tENT1*** | |
| GA_ENT1_22_23_GFP_FW | CAAATTCAAATGGTGAGCAAGGGC |
| GA_ENT1_L45G_RE01 | GATCCTTCGTGAGGATTGAGGCCAAGAGAAGTCTCCGGTCCG |
| GA_ENT1_L45G_FW01 | CGGACCGGAGACTTCTCTTGGCCTCAATCCTCACGAAGGATC |

| | |
|---|---|
| GA_ENT1_22_23_GFP_RE | GGTGGTCATCTTGTACAGCTCGTCCATG |
| **Cloning primers for introduction of *tENT1* into *NLS-eGFP-GUS* in *pENTR2B*** | |
| GA_NLS-GFP-GUS_backbone_FW01 | TCCTGTCCCAACAAAACCACCTCGAGATATCTAGACCCAGCTTTCTTGTAC AAAGTT |
| GA_NLS-GFP-GUS_backbone_RE01 | ATAACAAAGCTTTGGTAGAATCATTGTTTGCCTCCCTGCT |
| GA_ENT1_3UTR_NLS-GFP-GUS_FW01 | AGCAGGGAGGCAAACAATGATTCTACCAAAGCTTTGTTATCTGTATTTGT TGAA |
| GA_ENT1_3UTR_NLS-GFP-GUS_RE01 | CTGGGTCTAGATATCTCGAGGTGGTTTTGTTGGG |
| **Cloning primers for introduction of *pENT1* into *NLS-eGFP-GUS:tENT1* in *pENTR2B*** | |
| GA_ENT1_4kb_FW01 | CCGGAACCAATTCAGTCGACATGAACATATCA |
| GA_ENT1_wNterm_RE01 | CGTTTAAGAGAAGGCTGCATGGCTTCCGAGTCAGTTACGATTCCGG |
| GA_NLS-GFP-GUS_3UTR_backbone_FW01 | TCGTAACTGACTCGGAAGCCATGCAGCCTTCTCTTAAACGCA |
| GA_NLS-GFP-GUS_3UTR_backbone_RE01 | TATGGAGTTGATATGTTCATGTCGACTGAATTGGTTCCGG |
| **Cloning primers for CRISPR-Cas9 construct assembly** | |
| DT1-ENT1-3_F0 | TGACGATTCCGGCGGATTTATGTTTTAGAGCTAGAAATAGC |
| DT2-ENT1-3_R0 | AACACAACCAAGAACATCACAGCAATCTCTTAGTCGACTCTAC |
| DT1-ENT1-3_BsF | ATATATGGTCTCGATTGACGATTCCGGCGGATTTATGTT |
| DT2-ENT1-3_BsR | ATTATTGGTCTCGAAACACAACCAAGAACATCACAGC |
| **qPCR primers** | |
| qENT1_FW01 | CGTTGCCTTATCCGGTCTAGG |
| qENT1_RE01 | GTGACACAAGAACACCGGAGC |
| EF1α_F | TCCAGCTAAGGGTGCC |
| EF1α_R | GGTGGGTACTCGGAGA |
| Actin2_FW01 | AGACCTTTAACTCTCCCGCTATGTAT |
| Actin2_RE01 | GATTGGCACAGTGTGAGACACA |
| **Genotyping primers** | |
| LBb1.3 | ATTTTGCCGATTTCGGAAC |
| ENT1k_fwd | ATGACCACCACCGATAAATCC |
| ENT1l_fwd | ATGACTCCCATTAGTCAACGAATTT |
| ENT1_rev | TCAAATGACCCAGAACCAAGC |
| SALK_104866_LP | TGCAATGCAACTTGATAATGG |
| SALK_104866_RP | CACCATAACAACAATCCCCAC |
| SALK_025174_LP | ACCGTTGTCCATGATTTTCAG |
| SALK_025174_RP | GCGAGGCTCTTATGTGCATAG |

**Supplementary Table 2 | Primary root lengths of cytokinin-treated seedlings.** Median primary root lengths of the indicated experimental and reference lines grown on medium supplemented with DMSO, *t*ZR or iPR.

| **Experimental line** | **Reference line** | **Treatment** | **Experimental line median [cm]** | **Reference line median [cm]** |
|---|---|---|---|---|
| ENT1$^{WT}$-eGFP | Col-8 | DMSO | 7.68 | 7.19 |
| | | 100 nM *t*ZR | 5.57 | 6.16 |
| | | 100 nM iPR | 2.97 | 3.73 |
| ENT1$^{L45G}$-eGFP | Col-8 | DMSO | 6.96 | 6.54 |
| | | 100 nM *t*ZR | 3.55 | 5.45 |
| | | 100 nM *i*PR | 2.25 | 3.32 |
| *XVE>>ENT1$^{WT}$*, induced | *XVE>>ENT1$^{WT}$*, non-induced | DMSO | 2.17 | 7.33 |
| | | 100 nM *t*ZR | 0.87 | 6.37 |
| | | 100 nM iPR | 0.74 | 2.80 |
| *XVE>>ENT1$^{WT}$*-eGFP, induced | *XVE>>ENT1$^{WT}$*-eGFP, non-induced | DMSO | 4.58 | 7.62 |
| | | 100 nM *t*ZR | 1.17 | 5.78 |
| | | 100 nM iPR | 1.19 | 2.82 |
| *ent1-4* | Col-8 | DMSO | 7.42 | 7.10 |
| | | 100 nM *t*ZR | 5.89 | 5.87 |
| | | 100 nM iPR | 3.57 | 3.71 |
| ENT1$^{L45G}$-eGFP | Col-8 | DMSO | 7.22 | 6.88 |
| | | 50 nM *t*Z | 4.23 | 3.98 |
| | | 50 nM iP | 3.32 | 3.17 |

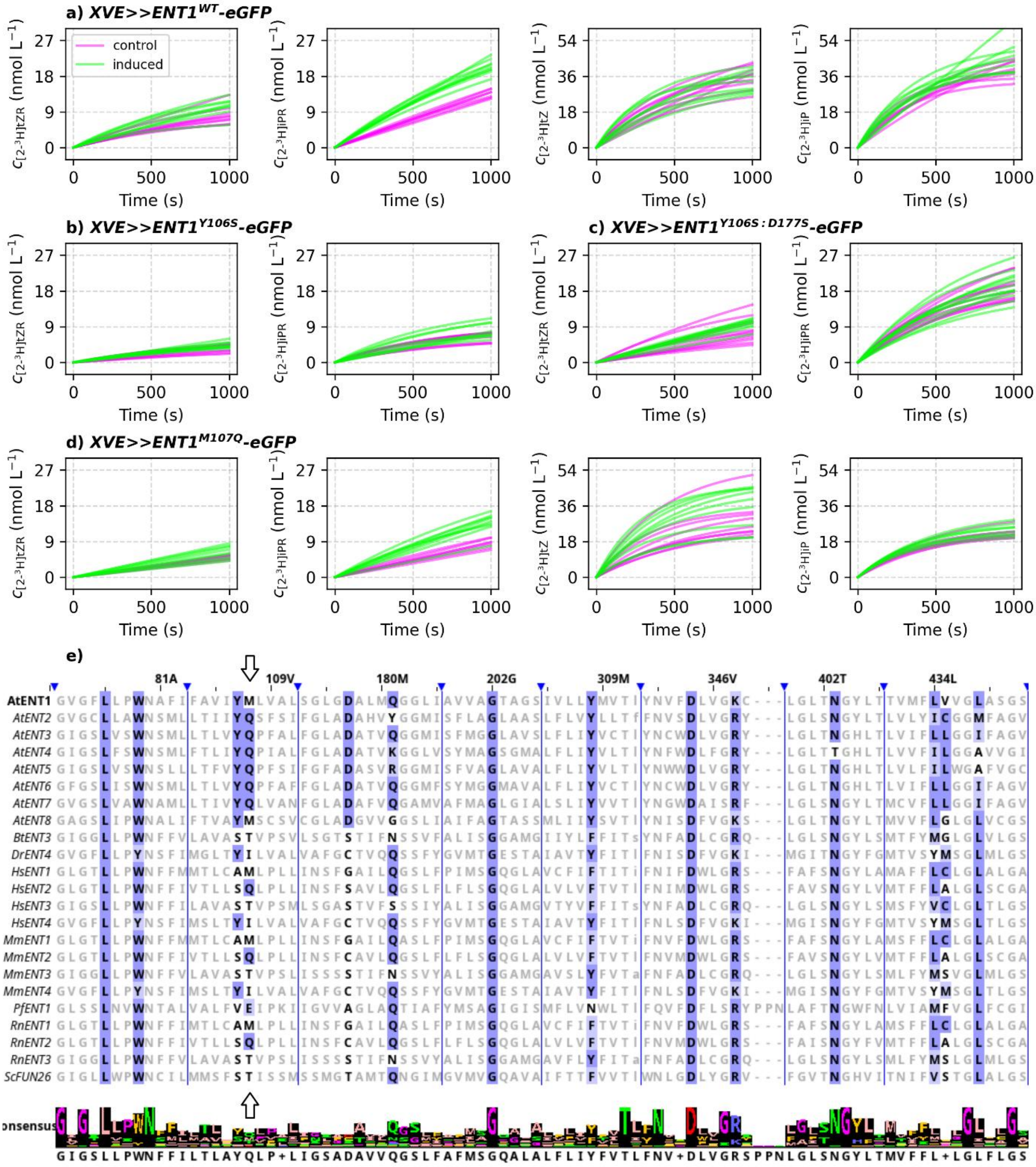


**Supplementary Fig. 1 | Time-course simulations of ENT1-mediated cytokinin uptake in BY-2 cell suspensions and sequence variability at the Met107-equivalent position in ENTs. a,** Simulated uptake of [2-$^3$H]*t*ZR, [2-$^3$H]iPR, [2-$^3$H]*t*Z and [2-$^3$H]iP in BY-2 cell suspensions carrying *XVE>>ENT1$^{WT}$-eGFP* construct from $t$ = 0 to $t$ = 1,000 s. **b-d,** As in **a**, but for BY-2 lines carrying *XVE>>ENT1$^{Y106S}$-eGFP*, *XVE>>ENT1$^{Y106S:D177S}$-eGFP* and *XVE>>ENT1$^{M107Q}$-eGFP* constructs, respectively. Concentrations, $c$, were calculated as functions of time according to equation (1). $L$ and $S$ were set to their optimized values, and $K$ was set to 0 to align the curves vertically. **e,** Alignment of ENT amino acid sequences from selected species. Highlighted positions correspond to ENT3 residues interacting with the docked pose of *t*Z; shading intensity denotes the BLOSUM62 score. Arrows indicate the positions corresponding to Gln62 in ENT3 and Met107 in ENT1. Vertical lines denote alignment breaks. Column annotations indicate the corresponding ENT1 residues. The alignment was visualized in Jalview. Species prefixes in the sequence labels denote *Arabidopsis thaliana* (*At*), *Bos taurus* (*Bt*), *Danio rerio* (*Dr*), *Homo sapiens* (*Hs*), *Mus musculus* (*Mm*), *Plasmodium falciparum* (*Pf*), *Rattus norvegicus* (*Rn*) and *Saccharomyces cerevisiae* (*Sc*). FUN26 denotes Function Unknown Now 26, the *S. cerevisiae* ENT homologue.

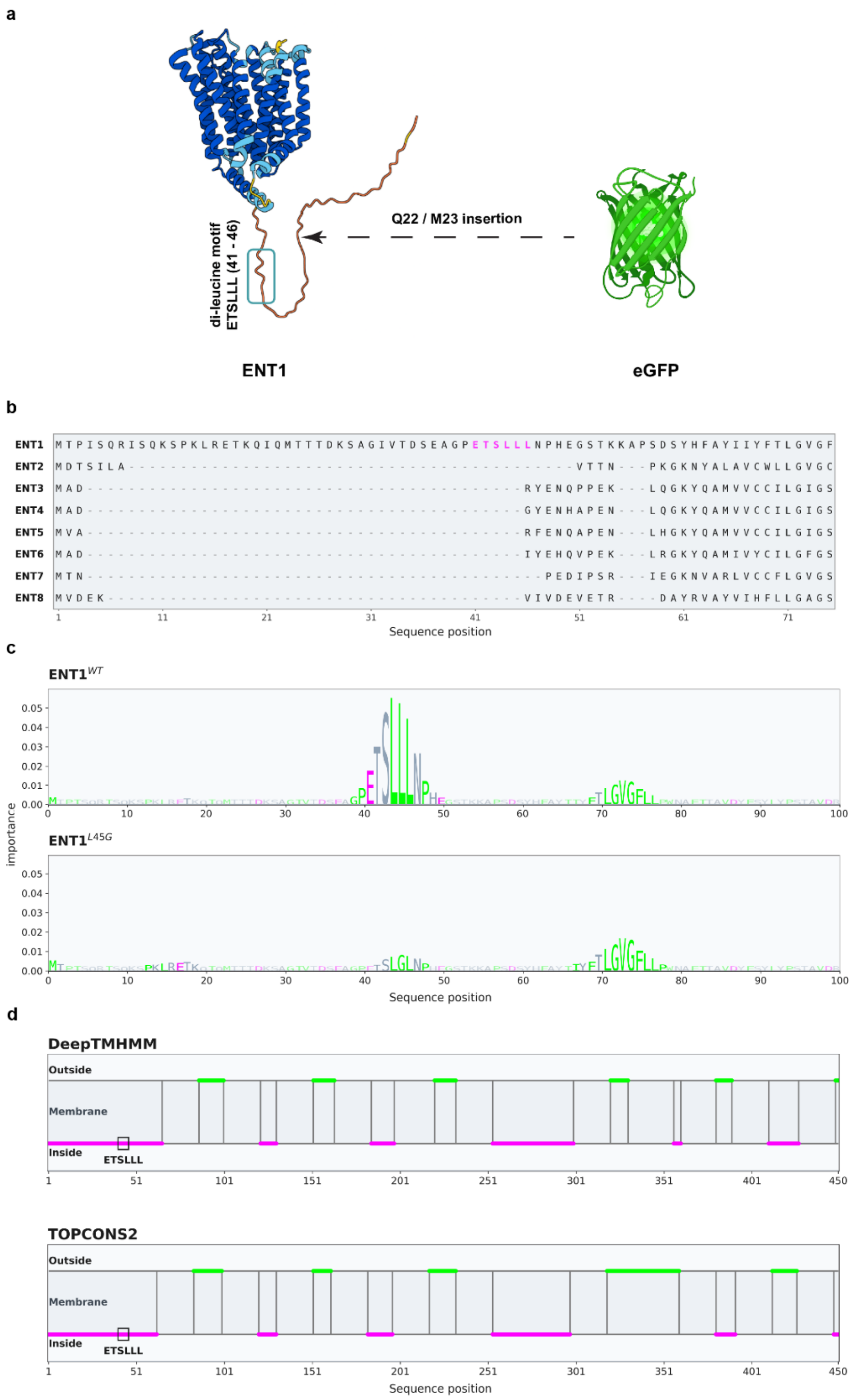


**Supplementary Fig. 2 | ENT1 internal eGFP tagging strategy and *in silico* support for a cytosolic N-terminal putative acidic di-leucine sorting motif. a,** AlphaFold DB v6 model of ENT1 (AF-Q8VXY7-F1, generated using the AlphaFold Monomer v2.0 pipeline) showing the internal eGFP insertion site between Gln22 and Met23. The N-terminal acidic di-leucine motif ETSLLL, located at residues 41-46, is highlighted with a blue box. **b,** N-terminal sequence alignment of Arabidopsis ENT transporters. The first 75 amino acids of ENT1-ENT8 are shown, with alignment gaps indicated by grey dashes. The dileucine-containing ETSLLL motif in ENT1 is highlighted in magenta. **c**, Residue-wise DeepLoc2.1 importance scores computed for full-length ENT1 and shown as an N-terminal zoom covering residues 1-100 for ENT1$^{WT}$ and ENT1$^{L45G}$. ENT1$^{WT}$ displays a pronounced importance peak around the ETSLLL-containing region, whereas the L45G substitution attenuates this signal. **d,** Predicted membrane topology of ENT1 from DeepTMHMM and TOPCONS2. In both predictions, the ETSLLL motif maps to a non-transmembrane N-terminal segment on the cytosolic side of the membrane.

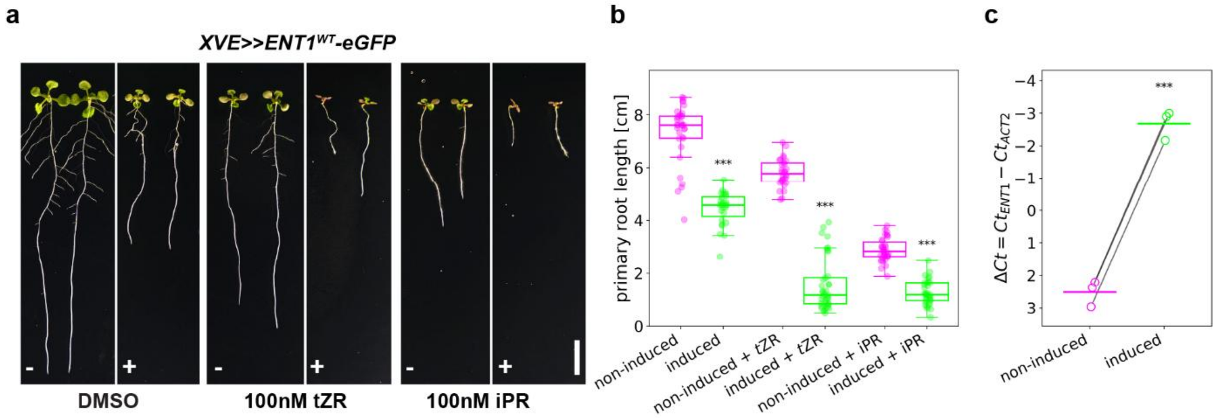


**Supplementary Fig. 3 | β-estradiol induction of ENT1$^{WT}$-eGFP enhances seedling sensitivity to cytokinin ribosides. a,** Representative phenotypes of 10-day-old *XVE>>ENT1$^{WT}$-eGFP* seedlings kept non-induced (-) or induced with 5 µM β-estradiol (+) on medium without cytokinins (DMSO) or supplemented with 100 nM *trans*-zeatin riboside (*t*ZR) or 100 nM isopentenyladenosine (iPR). **b,** Primary root length of seedlings shown in **a**. **c,** RT-qPCR analysis of *ENT1* transcript levels in *XVE>>ENT1$^{WT}$-eGFP* seedlings kept non-induced or induced with 5 µM β-estradiol. Expression is plotted as $\Delta C_t = C_{tENT1} - C_{tACT2}$; the y-axis is inverted, so lower $\Delta C_t$ values indicate higher expression. Open circles represent independent experiments ($n$ = 3), paired samples are connected by lines, and bars indicate means. Boxplots in **b** show the median and interquartile range (IQR); whiskers extend to the most extreme non-outlier values within 1.5 × IQR, and points indicate individual seedlings. Statistical significance was assessed using Welch's two-sample t-test (**b**) or a paired t-test (**c**). ns, not significant; $^{*}P < 0.05$; $^{**}P < 0.01$; $^{***}P < 0.001$. Scale bar: 1 cm.

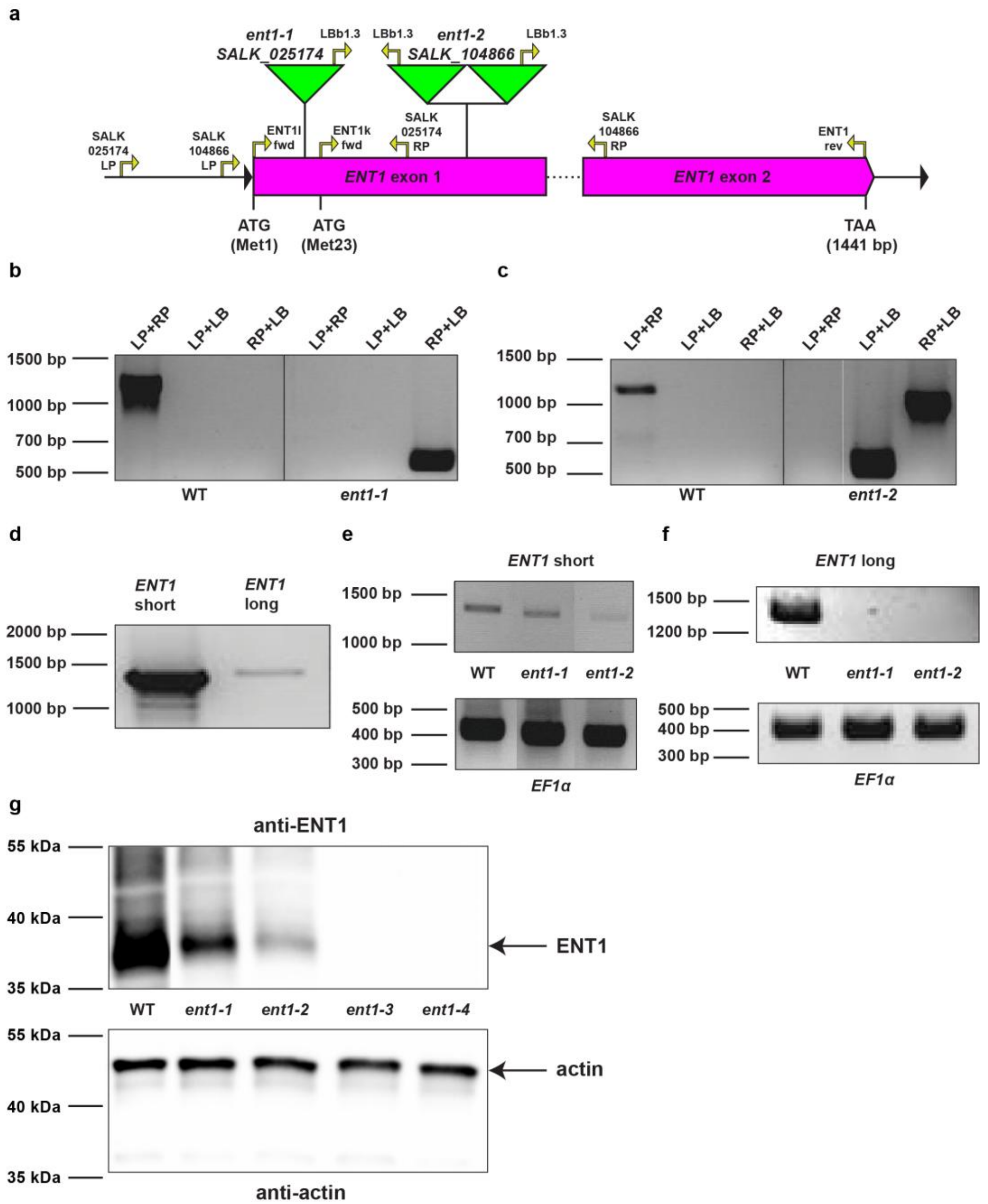


**Supplementary Fig. 4 | Genotyping and expression analysis of Arabidopsis *ent1* mutants. a,** Structure of the *ENT1* locus showing the positions of the T-DNA insertions in SALK_025174 (*ent1-1*) and SALK_104866 (*ent1-2*). Primer positions are indicated. **b,** PCR genotyping of genomic DNA from wild-type and *ent1-1*. The wild-type allele was amplified with the gene-specific primers SALK_025174_LP and SALK_025174_RP (1,117 bp). The T-DNA insertion was detected using the T-DNA left-border primer LBb1.3 in combination with SALK_025174_LP or SALK_025174_RP. **c,** PCR genotyping of genomic DNA from wild-type and *ent1-2*. The wild-type allele was amplified with the gene-specific primers SALK_104866_LP and SALK_104866_RP (1,119 bp). The T-DNA insertion was detected using LBb1.3 in combination with SALK_104866_LP or SALK_104866_RP. **d,** RT-PCR analysis of wild-type cDNA using primers amplifying the short *ENT1* transcript (ENT1k_fwd + ENT1_rev, 1,287 bp) or the long *ENT1* transcript (ENT1l_fwd + ENT1_rev, 1,353 bp). **e,** RT-PCR analysis of the short *ENT1* transcript in wild-type, *ent1-1* and *ent1-2* using ENT1k_fwd + ENT1_rev. *EF1α* was used as a control (Ef1α_fwd + Ef1α_rev, 404 bp). **f,** RT-PCR analysis of the long *ENT1* transcript in wild-type, *ent1-1* and *ent1-2* using ENT1l_fwd + ENT1_rev. *EF1α* was used as a control. **g,** Immunoblot analysis of ENT1 protein accumulation in 14-day-old seedlings of wild-type, *ent1-1*, *ent1-2*, *ent1-3* and *ent1-4* using anti-ENT1 antibody. A plant-specific anti-actin mouse monoclonal antibody was used as a loading control. The detection of actin and ENT1 proteins is denoted by black arrows.

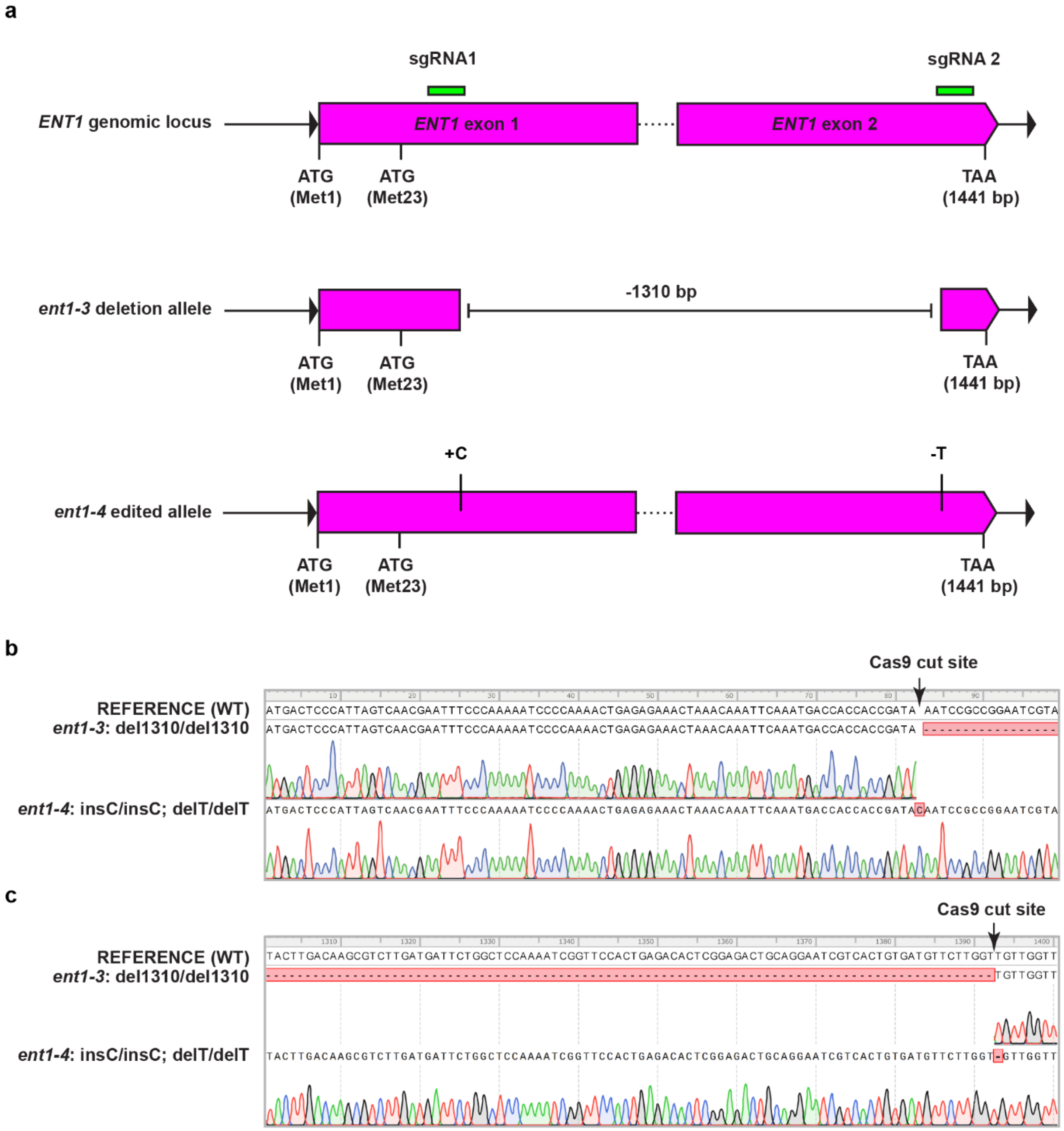


**Supplementary Fig. 5 | Generation and sequence validation of *ent1* CRISPR-Cas9 alleles. a**, Schematic representation of the *ENT1* genomic locus, sgRNA target sites and validated *ent1* mutant alleles. *ent1-3* contains a 1,310 bp deletion between the two sgRNA sites, whereas *ent1-4* retains the genomic structure but carries local indels at both target sites (+C and −T). **b,** Representative Sanger sequencing chromatogram of a PCR amplicon spanning the sgRNA1 cut site. **c,** As in **b**, but for the sgRNA2 cut site. Relative to the wild-type reference sequence, *ent1-3* knockout mutant carries a homozygous 1,310 bp deletion consistent with excision of the fragment between the two sgRNA cut sites, indicated by the dashed red region. *ent1-4* knockout mutant carries two homozygous 1-bp edits at the respective cut sites, a C insertion at sgRNA1 and a T deletion at sgRNA2, with the edited base indicated. Arrows mark the predicted Cas9 cut sites.

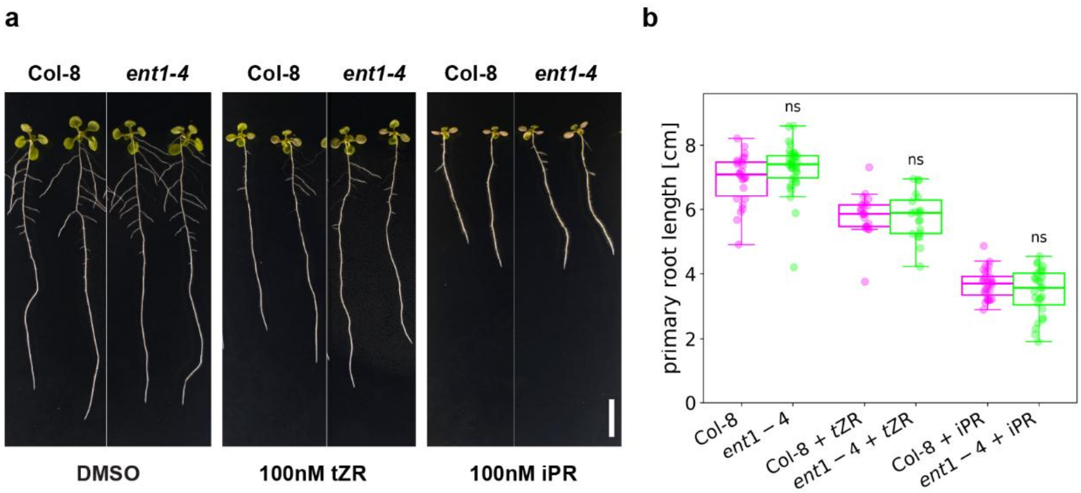


**Supplementary Fig. 6 | *ent1-4* seedlings show no increased sensitivity to cytokinin ribosides supplied in the growth medium. a,** Representative phenotypes of 10-day-old *A. thaliana* Col-8 and *ent1-4* seedlings grown on control medium containing DMSO or medium supplemented with 100 nM *trans*-zeatin riboside (*t*ZR) or 100 nM isopentenyladenosine (iPR). **b,** Primary root length for the genotypes and conditions shown in **a**. Boxplots show the median and interquartile range (IQR); whiskers extend to the most extreme non-outlier values within 1.5 × IQR, and points indicate individual seedlings. Statistical significance between Col-8 and *ent1-4* within each treatment was assessed using Welch's two-sample t-test; $^{*}P < 0.05$; ns, not significant. Scale bar: 1 cm.

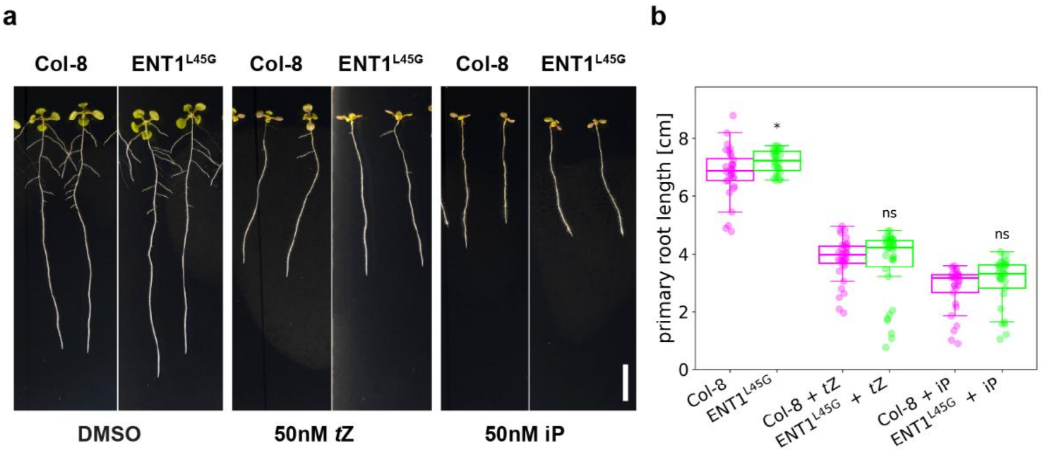


**Supplementary Fig. 7 | ENT1$^{L45G}$-eGFP seedlings show no increased sensitivity to cytokinin bases supplied in the growth medium. a,** Representative phenotypes of 10-day-old *A. thaliana* Col-8 and ENT1$^{L45G}$-eGFP seedlings grown on control medium containing DMSO or medium supplemented with 50 nM *trans*-zeatin (*t*Z) or 50 nM isopentenyladenine (iP). **b,** Primary root length for the genotypes and conditions shown in **a**. Boxplots show the median and interquartile range (IQR); whiskers extend to the most extreme non-outlier values within 1.5 × IQR, and points indicate individual seedlings. Statistical significance between Col-8 and ENT1$^{L45G}$-eGFP within each treatment was assessed using Welch's two-sample t-test; $^{*}P < 0.05$; ns, not significant. Scale bar: 1 cm.

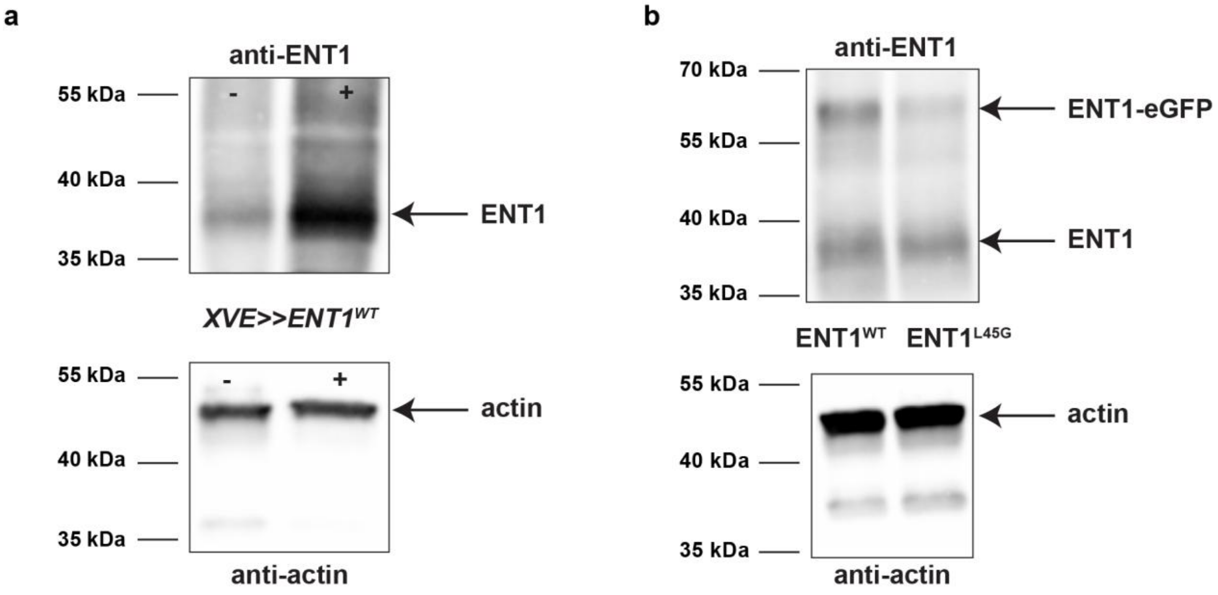


**Supplementary Fig. 8 | Immunoblot detection of ENT1 and ENT1-eGFP proteins in *XVE>>ENT1$^{WT}$*, ENT1$^{WT}$-eGFP and ENT1$^{L45G}$-eGFP lines. a,** Representative western blot of protein extracts from roots of 2-week-old inducible *XVE>>ENT1$^{WT}$* seedlings kept non-induced with DMSO (-) or induced with 5 µM β-estradiol (+). Immunoblotting was performed with an anti-ENT1 antibody; anti-actin antibody was used as a loading control. **b,** Representative western blot of protein extracts from roots of ENT1$^{WT}$-eGFP and ENT1$^{L45G}$-eGFP seedlings. The detection of actin, ENT1-eGFP fusion proteins and endogenous untagged ENT1 is denoted by black arrows. A plant-specific anti-actin mouse monoclonal antibody was used as a loading control. Mass spectrometry of the corresponding gel region confirmed the identity of the protein band.

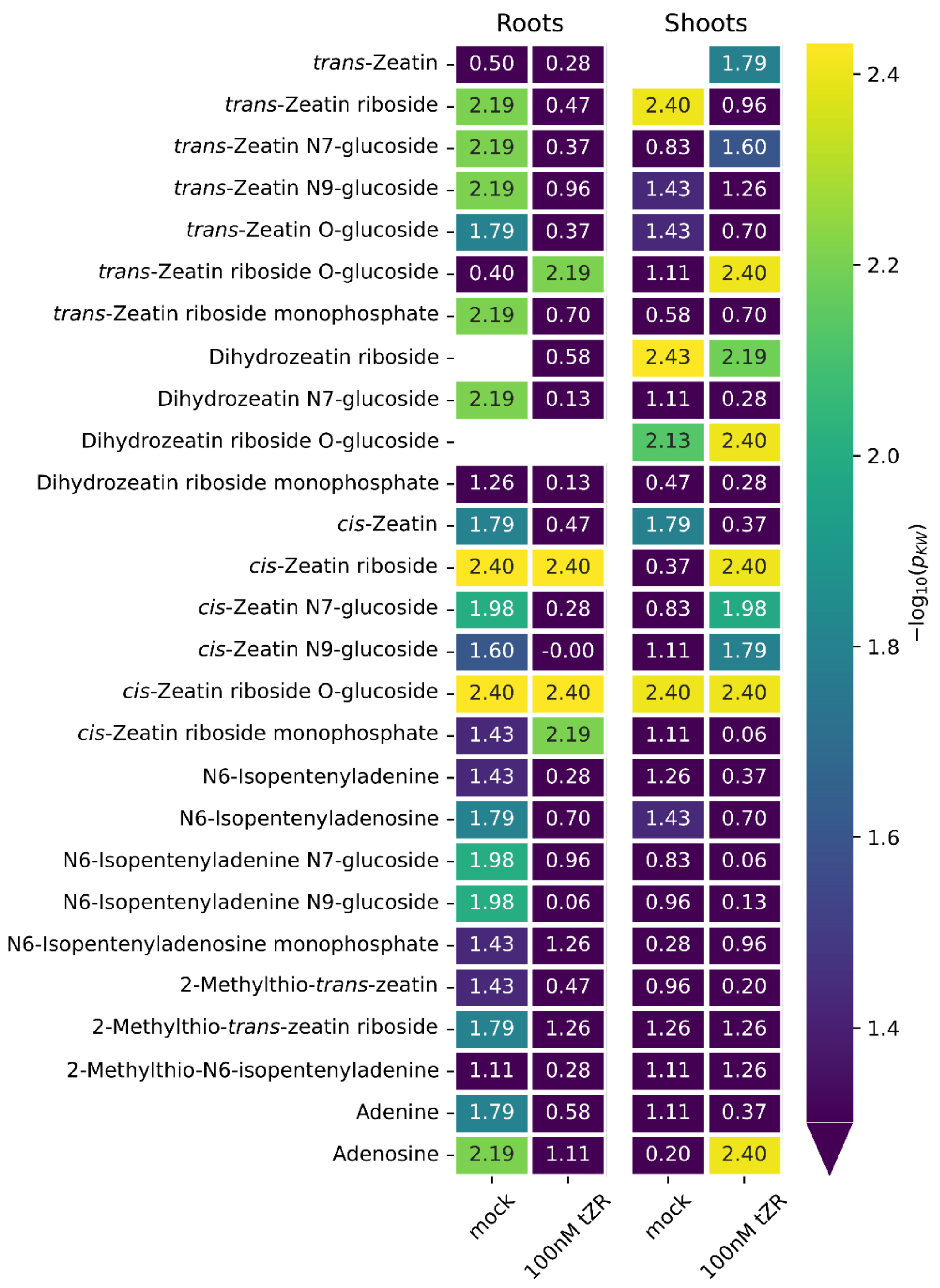


**Supplementary Fig. 9 | *P*-value map of genotype effects on individual cytokinin metabolites in Arabidopsis seedlings.** Heatmaps show $-\log_{10}$-transformed *P*-values obtained from Kruskal-Wallis tests comparing Col-8 and *ent1-4* roots and shoots treated with mock or 100 nM *t*ZR. Blank cells denote subsets in which all values were missing or equal to zero. The colour scale begins at $-\log_{10}(0.05)$, approximately 1.30.

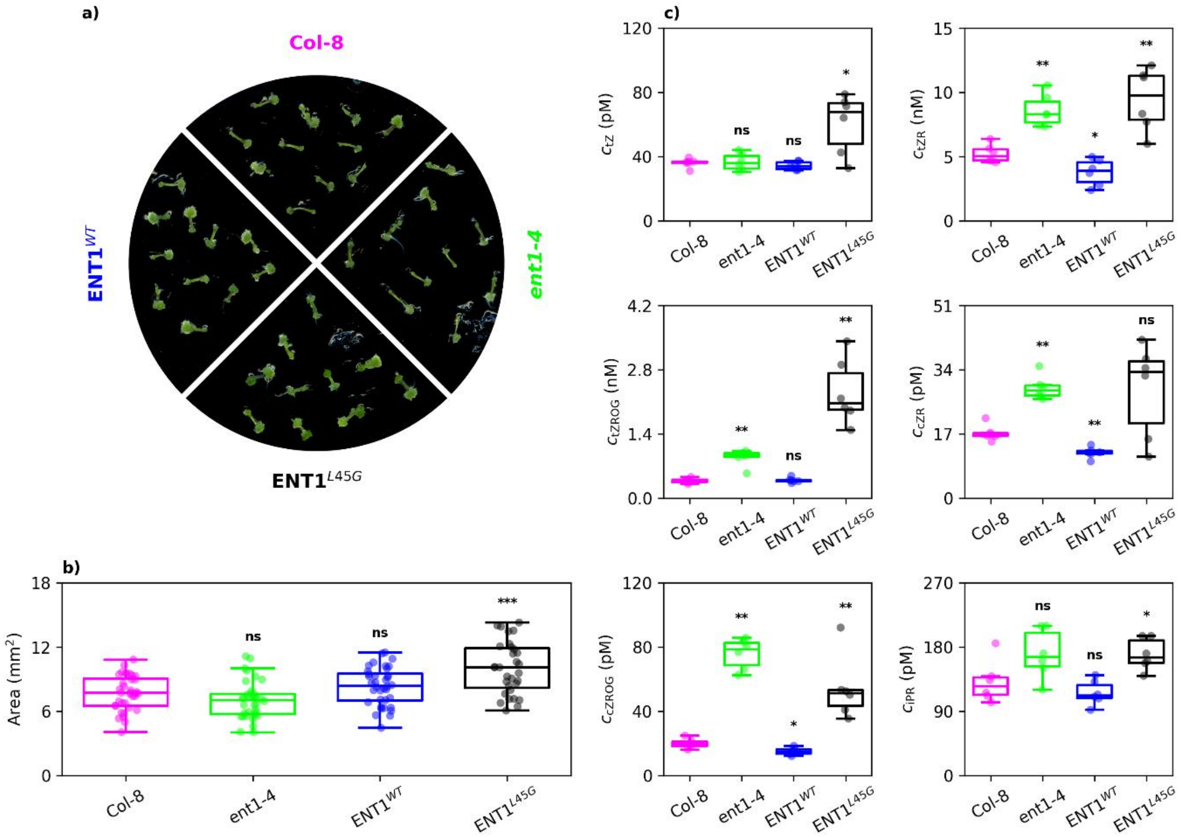


**Supplementary Fig. 10 | ENT1 modulates cytokinin riboside homeostasis and regeneration from Arabidopsis hypocotyl explants. a**, Representative image of hypocotyl explants of indicated genotypes after 21 days of regeneration on medium containing 100 µg $l^{-1}$ 1-naphthylacetic acid and 300 µg $l^{-1}$ *t*ZR. **b**, Area of regenerated hypocotyl explants. **c**, Metabolite profiles of CKs measured in tissues regenerated from the hypocotyl explants of the indicated genotypes. Boxplots show the median and interquartile range (IQR); whiskers extend to the most extreme non-outlier values within 1.5 × IQR, and points indicate individual measurements. Statistical significance between wild-type Col-8 and the indicated genotypes was assessed using a Kruskal-Wallis test; $^{*}P < 0.05$, $^{**}P < 0.01$, $^{***}P < 0.001$; ns, not significant.

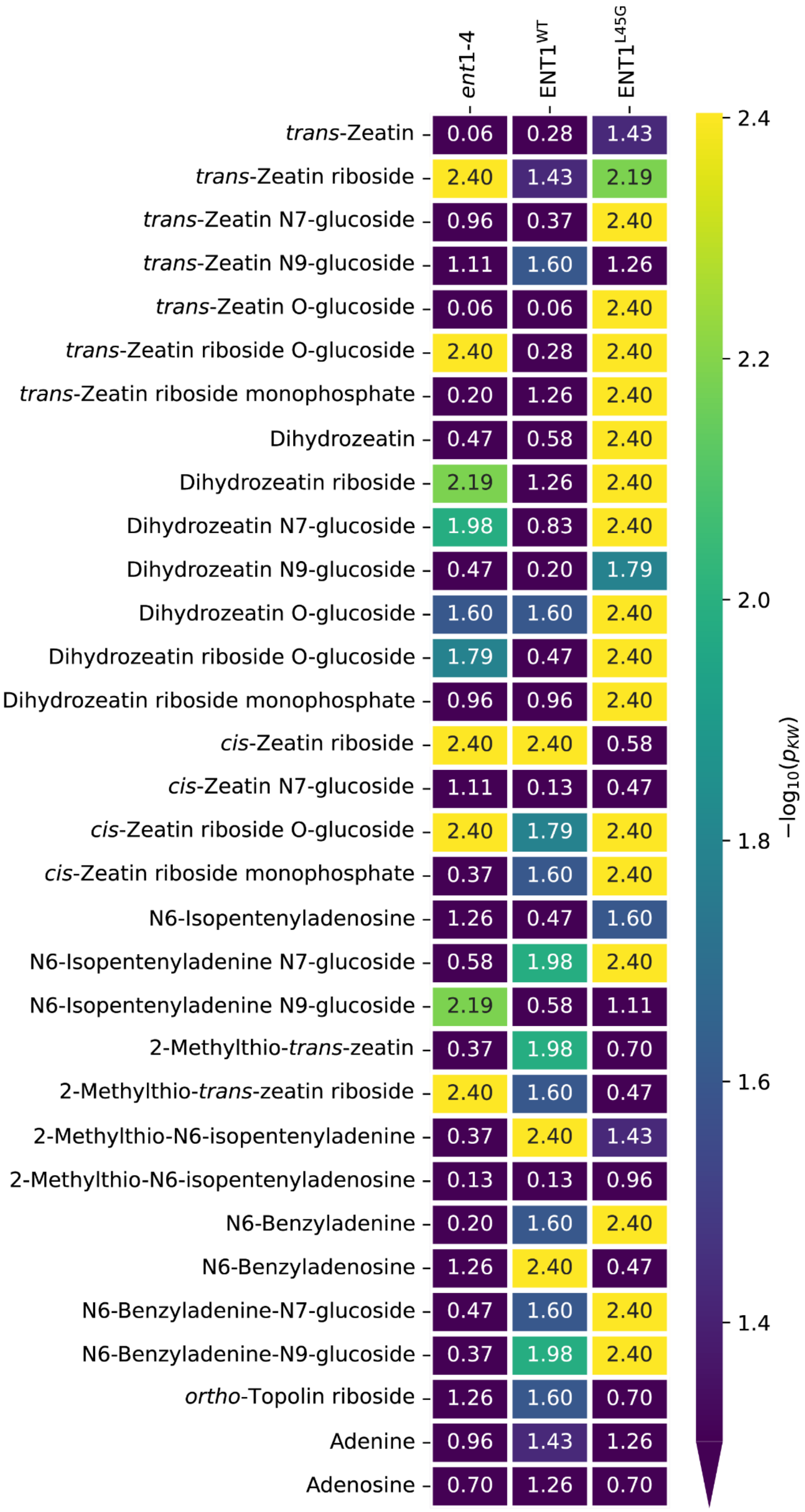


**Supplementary Fig. 11 | *P*-value map of genotype effects on individual cytokinin metabolites in Arabidopsis hypocotyl explants.** The heatmap shows $-\log_{10}$-transformed *P* values obtained from Kruskal-Wallis tests comparing Col-8 with *ent1-4*, ENT1$^{WT}$-eGFP, and ENT1$^{L45G}$-eGFP hypocotyl explants for each cytokinin species. The colour scale begins at $-\log_{10}(0.05)$, approximately 1.30.

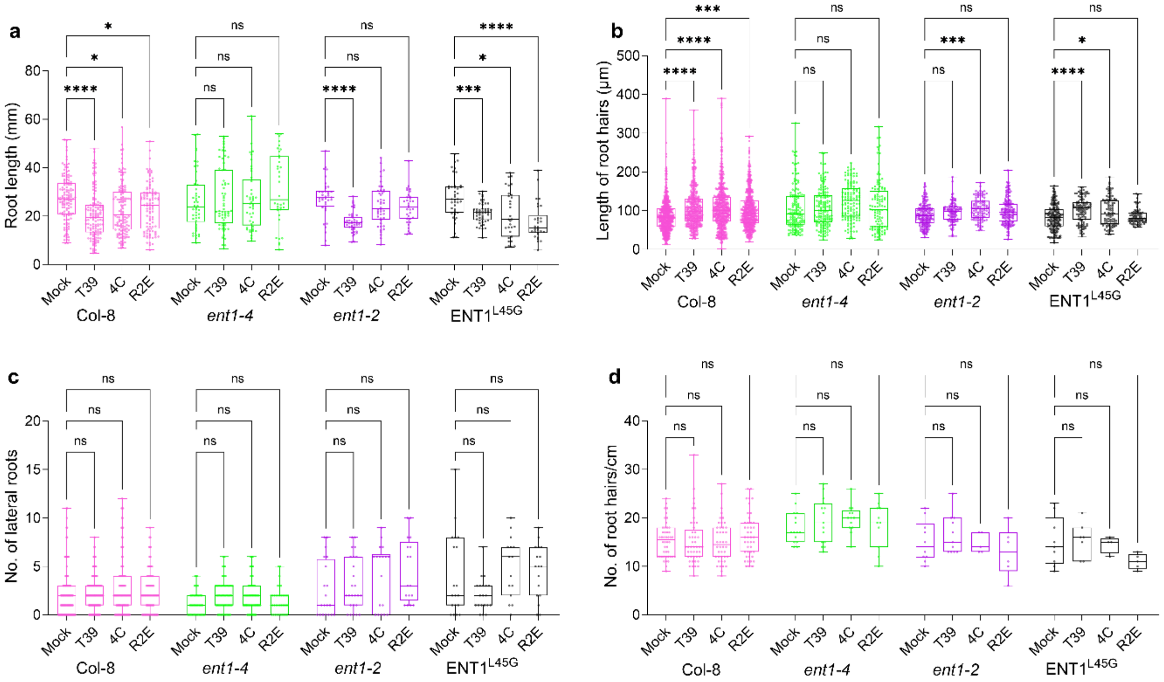


**Supplementary Fig. 12 | Root architectural responses to cytokinin-producing microbes are altered in lines with altered ENT1 function.** *Arabidopsis thaliana* seeds of the indicated genotypes were germinated on vertical agar plates and treated in vitro with the indicated microbes 4 days after germination. Root architecture parameters were examined after 4 days of co-cultivation: **a,** root length. **b**, root hair length. **c**, number of lateral roots. **d,** number of root hairs in the 1-cm region above the meristematic zone. Three independent experiments were conducted. Boxplots show the median and interquartile range (IQR); whiskers extend from minimum to maximum values, and all individual data points are shown. Asterisks indicate statistically significant differences within each genotype relative to mock treatment, assessed using Welch's ANOVA followed by Dunnett's post hoc test. **a,** $n > 30$; **b,** $n > 100$; **c,** $n > 15$; **d,** $n > 6$. ns, not significant; $^{*}P < 0.05$, $^{**}P < 0.01$, $^{***}P < 0.001$, $^{****}P < 0.0001$.
